# eQTL in diseased colon tissue identifies novel target genes associated with IBD

**DOI:** 10.1101/2024.10.14.618229

**Authors:** Nina C. Nishiyama, Sophie Silverstein, Kimberly Darlington, Meaghan M. Kennedy Ng, Katelyn M. Clough, Mikaela Bauer, Caroline Beasley, Akshatha Bharadwaj, Rajee Ganesan, Muneera R. Kapadia, Gwen Lau, Grace Lian, Reza Rahbar, Timothy S. Sadiq, Matthew R. Schaner, Jonathan Stem, Jessica Friton, William A. Faubion, Shehzad Z. Sheikh, Terrence S. Furey

## Abstract

Genome-wide association studies (GWAS) have identified over 300 loci associated with the inflammatory bowel diseases (IBD), but putative causal genes for most are unknown. We conducted the largest disease-focused expression quantitative trait loci (eQTL) analysis using colon tissue from 252 IBD patients to determine genetic effects on gene expression and potential contribution to IBD. Combined with two non-IBD colon eQTL studies, we identified 194 potential target genes for 108 GWAS loci. eQTL in IBD tissue were enriched for IBD GWAS loci colocalizations, provided novel evidence for IBD-associated genes such as *ABO* and *TNFRSF14*, and identified additional target genes compared to non-IBD tissue eQTL. IBD-associated eQTL unique to diseased tissue had distinct regulatory and functional characteristics with increased effect sizes. Together, these highlight the importance of eQTL studies in diseased tissue for understanding functional consequences of genetic variants, and elucidating molecular mechanisms and regulation of key genes involved in IBD.

## Introduction

The inflammatory bowel diseases (IBD), namely Crohn’s disease (CD) and ulcerative colitis (UC), are complex, chronic inflammatory diseases of the gastrointestinal (GI) tract^1–4^. The most recent meta-analysis of genome-wide association studies (GWAS) identified 320 loci associated with IBD^5^. Despite this, little is known about how most of these genetic variants functionally contribute to disease. Like other GWAS traits, most IBD-associated variants fall within non-coding genomic regions, making it challenging to determine their direct functional consequences^6^. Mapping expression quantitative trait loci (eQTL) in trait-relevant tissue allows us to associate genetic variants with expression of nearby genes^7–9^. Furthermore, colocalizing eQTL and GWAS variants has successfully aided in the interpretation of GWAS loci through target gene identification, with over 40% of GWAS loci explained by eQTL colocalizations for some traits and tissues, such as the role of adipose gene expression in cardiometabolic traits^10,11^. However, current eQTL studies, such as from the GTEx Consortium and the BarcUVa-Seq project, only explain ∼20% of IBD-associated loci using steady-state colon tissue^12,13^. One possible explanation for this limited overlap is that GWAS hits may only colocalize with eQTL determined in the proper context such as key cell types, in response to specific stimuli, or at critical time points^14,15^. Indeed, the addition of context-specific eQTL can explain additional GWAS loci by as much as 30% that are missed by “baseline” or normal tissue eQTL^16^.

Studies only using healthy donors may fail to capture the range of eQTL effects found in disease states. Other studies have identified disease-interacting and specific eQTL that point towards cell-type-specific mechanisms underlying disease biology, are enriched for disease-relevant transcription factor binding motifs, and have identified novel genes key to disease progression^17,18^. However, few studies have focused solely on mapping eQTL in the context of disease. Hu et al. performed an intestinal eQTL meta-analysis that combined both colonic and ileal samples from a cohort of 171 IBD patients, with a focus on inflammation-dependent intestinal eQTL. While they showed that inflammation status can reveal the regulatory effects for some variants, they did not report on novel target genes in GWAS loci using diseased tissue. Furthermore, other colon-specific eQTL studies in IBD tissue have been limited either in sample size, sequencing depth, sampling location, or have included other factors which may introduce additional variation^19–23^. Finally, time and cost burdens have been limiting factors to conducting diseased-focused eQTL studies on a scale to match with non-diseased eQTL references such as GTEx.

To address this, we have performed the largest, single tissue disease-focused eQTL analysis using colon tissue samples from 252 IBD patients. We compared our results to published non-IBD colon eQTL studies and colocalized eQTL from both IBD-focused and non-IBD studies with IBD GWAS loci to identify potential target genes. Across the three studies, we identified 194 potential target genes with a diverse range of biological functions relevant to IBD biology. We are the first to show evidence for 49 potential target genes in colon for 29 of the 81 newly reported loci by Liu et al., including robust evidence for 11 genes identified using at least two independent cohorts^5^. We found that eQTL from our IBD-focused study were more strongly enriched for GWAS colocalizations compared to the non-IBD studies. Most importantly, we found that mapping eQTL in diseased tissue revealed novel colocalizations and a unique set of potential target genes not uncovered using the larger non-diseased cohorts. This set included evidence for the first genetic links to *ABO* and *TNFRSF14* expression in the colon, both of which have previously been shown to contribute to changes in the intestinal environment also observed in IBD^24–28^. In some loci, such as for *TNFRSF14,* different eQTL and target genes colocalized in diseased and non-diseased tissue. We discovered distinct regulatory characteristics of our novel eQTL-GWAS colocalizations in contrast to findings using non-IBD tissue. Disease-associated eQTL from IBD tissue tended to be more distal to the target gene and were associated with different classes of predicted biological function. Some genes have larger eQTL effect sizes in IBD tissue compared to in non-IBD tissue, suggesting the genetic effect may be amplified in the presence of disease. These results suggest eQTL mapping in disease tissue is critical in comprehensively identifying and understanding regulatory effects of variants and genes within GWAS loci.

## Results

### Unique eQTL mapped in IBD tissue suggest novel regulatory activity in the presence of disease

We mapped *cis*-eQTL in colon tissue from 252 IBD patients, which we refer to as UNC eQTL, using 30,715 genes and 8.4M variants (MAF ≥ 0.02). We identified 457,397 significant *cis*-eQTL (FDR < 0.05) associated with 4,087 genes (eGenes, **Supplementary Table 1**). We detected at least two independent signals for 268 eGenes using conditional mapping (**Supplementary Table 2**). Like other studies, we found the majority of primary lead eQTL variants (eVariants) to be located proximal to the transcription start site (TSS) of the target eGene with a median absolute distance of 23.95 kb (**Supplementary Figure 1**). Similarly, secondary and tertiary lead eVariants tended to be more distal to target eGene TSSs compared to primary lead eVariants (**Supplementary Figure 2**). Given our limited power to detect secondary and higher order signals, all subsequent analyses were focused on primary signals only.

Next, we compared our UNC eQTL to eQTL mapped in non-diseased colon tissue (non-IBD eQTL) using publicly available summary statistics from GTEx (v8, GTEx eQTL) and BarcUVa-Seq (BarcUVa eQTL, **Supplementary Table 3**)^12,13^. We found that 15% of lead eVariant-eGene pairs in UNC eQTL were also lead pairs in GTEx and/or BarcUVa eQTL. Effect size estimates for these shared lead eVariant-eGene pairs were correlated between UNC and non-IBD eQTL (Pearson’s r = 0.922, p = 6.14e-172 for GTEx; r = 0.958, p = 4.83e-189 for BarcUVa) and were comparable to the effect size correlation between GTEx and BarcUVa eQTL (r = 0.900, p = 4.05e-191; **Figure 1a**). Effect size estimates remained correlated when considering all significant shared eVariant-eGene pairs: UNC vs. GTEx eQTL r = 0.865; UNC vs. BarcUVa eQTL r = 0.898; GTEx vs. BarcUVa eQTL r = 0.882 (**Supplementary Figure 3**).

**Figure 1.**
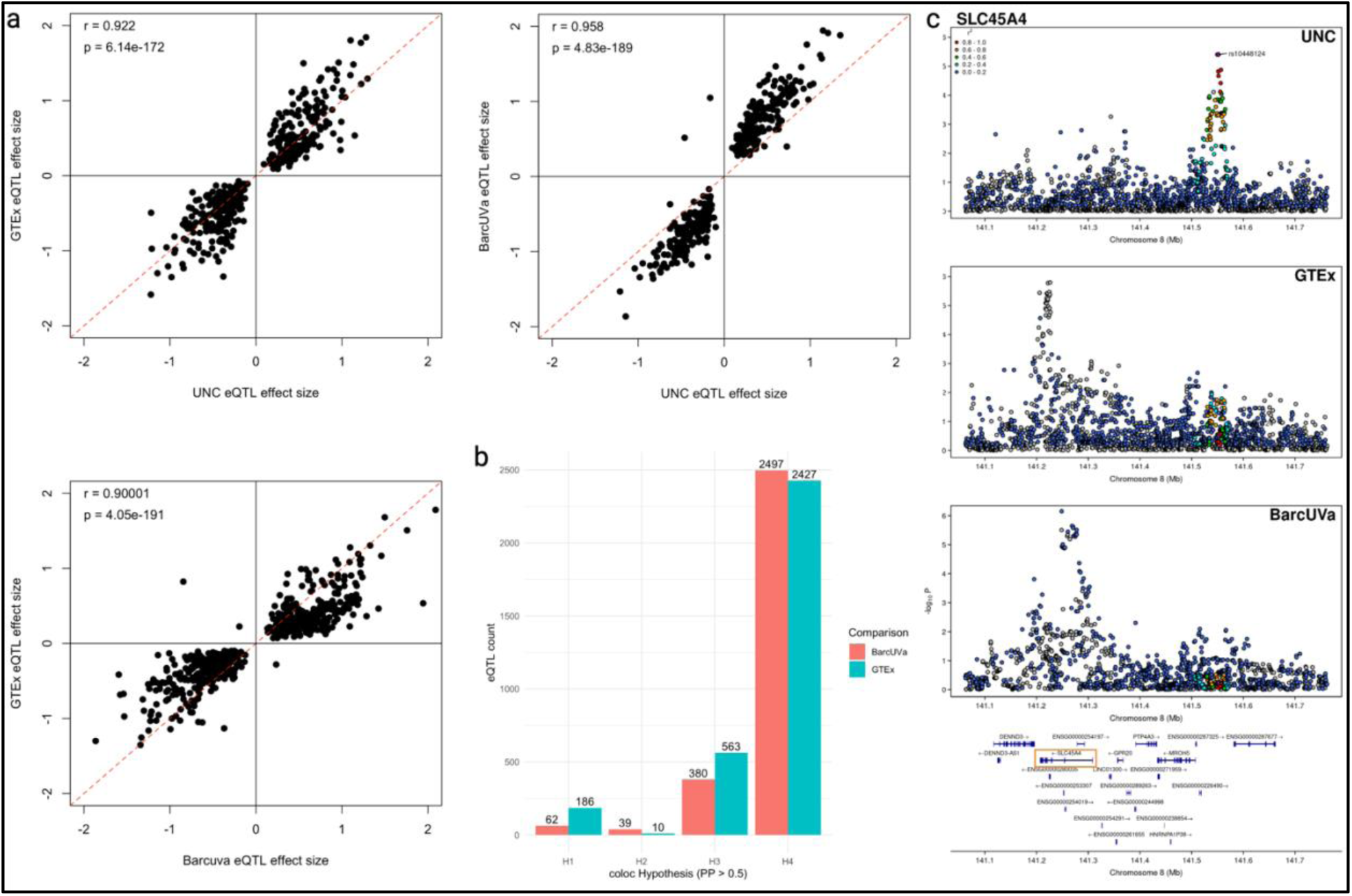
Colon eQTL are robust across disease states. a. Pairwise scatter plots of effect size estimates for shared primary lead eVariant-eGene pairs are highly correlated across studies. Pearson’s correlation coefficient and p-values for effect size estimates are shown. Red dashed line represents the (0,1) intercept. b. Bar plot of UNC eQTL-non-IBD eQTL colocalization results by coloc hypothesis (PP > 0.5): H1, significant signal detected in UNC data only; H2, significant signal detected in non-IBD data only; H3, significant signal detected in both data sets but different causal variants; H4, significant signal detected in both data sets, same causal variant. c. Stacked LocusZoom plots for SLC45A4 eQTL signals across disease states in the colon suggest a unique genetic association in IBD tissue. Index variant for the UNC eQTL is labeled with a purple diamond.

To comprehensively determine shared signals across studies, we performed colocalization on eQTL mapped in at least one study in 15,337 commonly tested genes (**Supplementary Tables 4-6**). For UNC eQTL, 3468 (84.85%) eGenes were also tested in the GTEx and/or BarcUVa eQTL analyses. Of these, 3050 UNC eQTL (87.94% tested) colocalized with non-IBD colon eQTL (*coloc* PP4 > 0.5), suggesting that the majority of eQTL signals are shared across disease states within the colon (Figure **1b**). Interestingly, 157 (4.53%) UNC eQTL did not colocalize with GTEx nor BarcUVa eQTL (*coloc* PP3 > 0.5 for both comparisons), and an additional 153 eQTL (4.41%) were supported by a PP3 > 0.5 in either GTEx or BarcUVa, suggesting that some eQTL may be driven by regulatory mechanisms only active or more active in the presence of disease (**Figure 1c**). An analogous analysis to colocalize non-IBD eQTL revealed that 12.03% of GTEx eQTL and 12.20% of BarcUVa eQTL did not colocalize with each other (PP3 > 0.5; **Supplementary Figure 4**).

To explore the robustness of our UNC eQTL, we calculated the replication rate of the p-value distribution (Storey’s π1) for UNC eQTL in the GTEx and BarcUVa data sets. UNC eQTL showed high replication in both GTEx eQTL (π1 = 0.912) and BarcUVa eQTL (π1 = 0.949). These values are higher than replication rates reported between GTEx and BarcUVa (π1 = 0.76)^13^.

In summary, the majority of the eQTL signals we observed in IBD colon tissue are shared and highly replicable with non-IBD colon eQTL, demonstrated with two external and independent cohorts. Furthermore, effect sizes for shared eVariant-eGene pairs were well-correlated across data sets. Together these results suggest that most colon eQTL are robust across disease states. However, we do detect some UNC eQTL signals that do not seem to be observed or shared with non-IBD eQTL, even with larger samples sizes. This suggests unique regulatory activity in the presence of disease uncovers alternate eQTL which may reveal new insights into IBD-associated loci beyond our present knowledge using only non-IBD eQTL.

### eQTL in IBD tissue are strongly enriched for IBD GWAS variants

To assess the degree of genetic overlap with IBD, we quantified the enrichment of eVariants in IBD GWAS loci compared to all variants tested for eQTL. We surveyed a sampling of tissues with both known and unlikely involvement in the disease, comparing to data primarily generated by GTEx^29^. The UNC eQTL had the strongest enrichment across CD-, UC-, and IBD-associated variants, while BarcUVa eQTL had the weakest enrichment across all three disease traits (**Figure 2a**). Among other GTEx tissues we tested, adipose eQTL were the least enriched for disease-associated variants. Interestingly, when we compared the enrichment scores for UNC eQTL vs. GTEx transverse colon and BarcUVa eQTL, we observed a substantial increase in the enrichment scores for CD (Pearson’s χ^2^ = 6554.2, df = 2, p < 2.2e-16), UC (Pearson’s χ^2^ = 6787.5, df = 2, p < 2.2e-16), and IBD (Pearson’s χ^2^ = 6964.9, df = 2, p < 2.2e-16) using IBD tissue, suggesting the increased overlap in the presence of IBD-associated variants among UNC eVariants may indicate a greater degree of shared genetic architecture.

**Figure 2.**
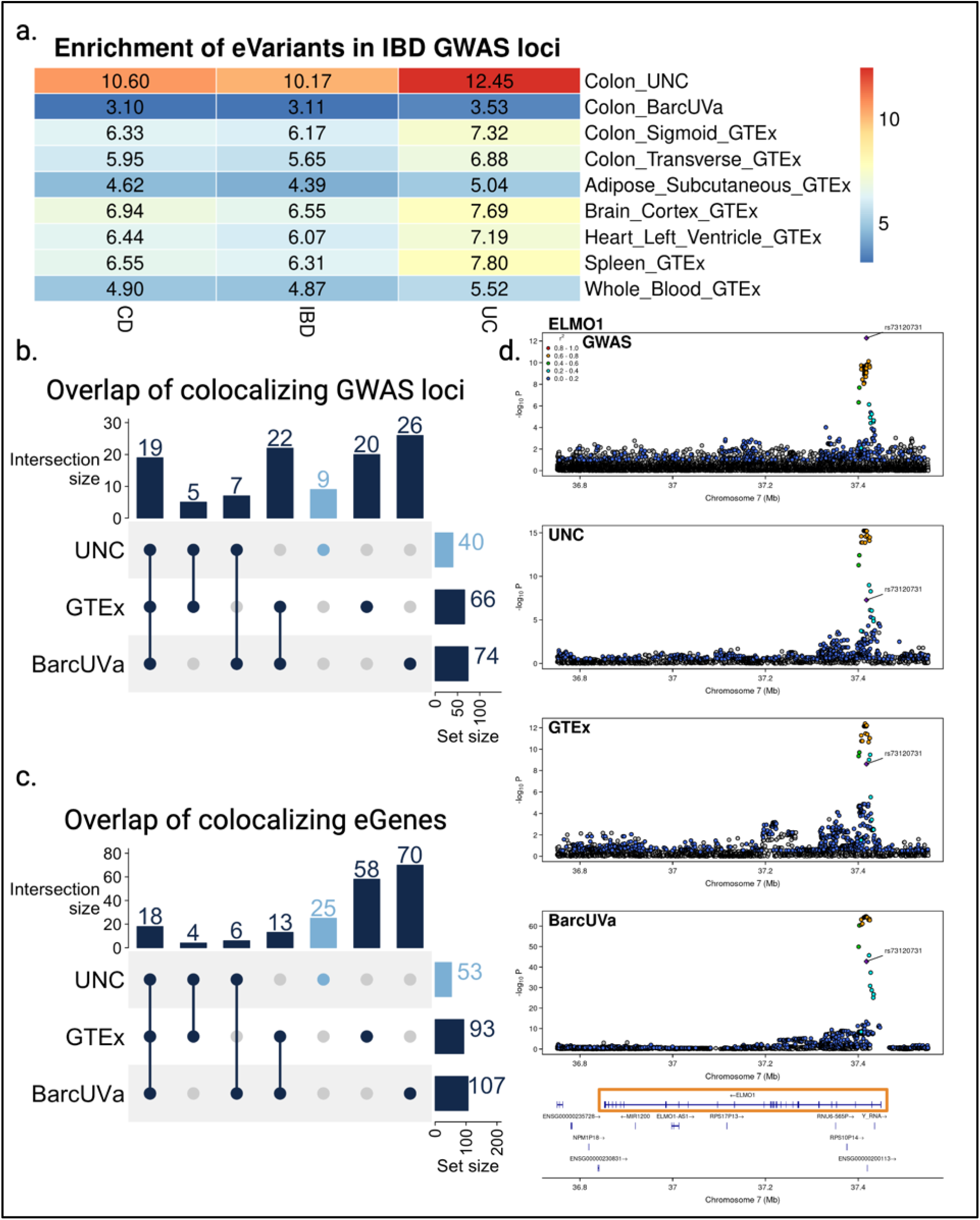
GWAS loci colocalize with eQTL across disease states. a. Heatmap of enrichment fold change scores of eVariants among IBD GWAS variants showed increased enrichment in eQTL from IBD colon tissue. b. UpSet plot of the overlap GWAS loci that colocalize with colon eQTL, with those unique to IBD tissue in Carolina blue. c. UpSet plot of overlapping colon eGenes that colocalize with GWAS loci, with those unique to IBD tissue in Carolina blue. d. Stacked LocusZoom plot for the colocalizing GWAS and ELMO1 eQTL signal shows consistent genetic associations found across diseases states in the colon. The CD GWAS index variant is labeled with a purple diamond.

### IBD-linked variants regulate colonic genes associated with diverse functional roles relevant to disease biology

We colocalized colon eQTL from all three data sets with 320 IBD-associated loci to identify shared disease-relevant genetic signals and putative target genes^5^. Out of 720 eQTL-GWAS pairs tested, 240 eQTL signals, associated with 194 unique genes, colocalized with 108 GWAS loci in at least one data set (PP4 > 0.5), increasing the proportion of IBD loci explained by colon tissue eQTL to 34% (**Figure 2b**; **Supplementary Tables 7-9**). Over 20% of these eGenes colocalized with GWAS in at least two eQTL data sets, including 18 that colocalized in all three (**Figure 2c**). These included known genes involved in regulating mucosal immune responses such as *FUT2*, *ERAP2*, *IRF5*, *CXCL5*, *LTBR*, and *CTSW*^26,30–40^.

The most recent meta-analysis from the International IBD Genetics Consortium (IIBDGC) has identified 81 new IBD-associated loci^5^. Across all three colon eQTL data sets, we found colon eQTL for 49 eGenes that colocalized with 29 out of the 81 newly reported GWAS loci (**Supplementary Figure 5**). This goes beyond the nine target genes identified by Liu et al. for the nine GWAS index variants that overlapped fine-mapped GTEx eQTL in any tissue, none of which were found to overlap eQTL in colonic mucosa. Furthermore, our results provide parallel evidence of disease-associated regulatory effects in the colon for five of these genes. We found that eQTL for 11 of 49 eGenes colocalized with ten of these new loci in at least two colon eQTL data sets, with six eGenes colocalizing in all three cohorts: *TMEM170A, HORMAD1, CDK18, DR1, FLRT3,* and *ELMO1* (**Figure 2c, 2d**). Thus, shared results across multiple independent cohorts can provide robust target gene predictions for recently reported loci. Both *FLRT3* and *ELMO1* are suggested to be involved in host responses to E. coli^41,42^. Interestingly, *ELMO1* has been reported to interact with *NOD2*, a well-known intracellular innate immune cell microbial sensor with polymorphisms linked to CD risk^43,44^. These results could suggest a potential genetic interaction between *NOD2* and *ELMO1* variants that may influence responses to enteric pathogens in some individuals.

Next, we queried gene ontology (GO) for biological process terms associated with colocalizing eGenes to survey the plausible higher order functional impact of the potential target genes contributing to IBD. One hundred twenty-eight colocalizing eGenes were found to be associated with a total of 936 unique GO terms, ranging from 1-94 GO terms/eGene, and with a median of six GO terms/eGene (**Supplementary Figure 6**; **Supplementary Table 10**). Using semantic similarity scores and hierarchical clustering, we clustered GO terms into six primary clusters and 54 sub-clusters, ranging from 4-16 sub-clusters under a given primary cluster, and a seventh category was included for the remaining colocalizing eGenes without annotated GO terms (**Supplementary Figure 7**). The primary clusters were represented by the following broad categories: cell adhesion, cell differentiation, immune response, cell proliferation, transcriptional regulation, and signal transduction. Across all three eQTL studies, the largest proportion of eGenes mapped to transcriptional regulation. In some cases, we found that eGenes associated with multiple terms mapped to multiple primary clusters (median = 2 primary clusters/eGene). Multifunctional eGenes may be indicative of key regulators of disease, such as *SMAD4* and *PARK7*^45–48^. Collectively, these results highlight the diverse function of genes potentially contributing to IBD biology.

### Mapping eQTL in diseased tissue reveals novel GWAS colocalizations for genes with disease-relevant functions not found in non-IBD tissue

We compared our UNC eQTL colocalization results to the GTEx and BarcUVa eQTL colocalization results to determine whether disease state affected GWAS colocalization discovery. Interestingly, we found that most GWAS colocalizations were unique to each study. Despite this, we observed that our UNC eQTL were 1.5x more likely to colocalize with IBD GWAS loci compared to non-IBD eQTL (Pearson’s χ^2^ = 8.31, df = 2, p = 0.0157). Using IBD tissue, we identified a unique set of 25 eQTL that colocalized with 20 GWAS loci (**Figure 2c**). We employed a cross-colocalization strategy to determine whether these signals were also detected in colon tissue in the absence of disease. When eGenes associated with colocalizing UNC eQTL signals were tested in the GTEx and BarcUVa data, they did not colocalize with GWAS loci, despite significant eQTL reported by GTEx and/or BarcUVa for 18 out of these 25 eGenes. Furthermore, only four of the 25 uniquely colocalizing UNC eGenes have been previously reported to be associated with IBD using intestinal data from a separate, smaller IBD cohort^23^. Finally, using the UNC IBD tissue data set we report novel colocalizations in colon tissue for nine GWAS loci, implicating 12 potential novel target genes discovered only by using diseased tissue (**Figure 2b**; **Supplementary Table 7**).

In contrast, when we considered the complementary results for the non-IBD eQTL that uniquely colocalize with GWAS, we did not observe the same patterns of exclusivity that we observed with our UNC eQTL findings. Using our cross-colocalization strategy, we tested the set of 141 eGenes found to colocalize with non-IBD eQTL (**Figure 2c**). Of these, we found we were able to “rescue” additional GWAS colocalizations for 15 UNC eQTL (PP4 > 0.5) that were not originally tested for colocalization. This number increased to 21 UNC eQTL when we consider the *coloc* hypothesis with the highest posterior probability (0.439 ≤ PP4 ≤ 0.961; **Supplementary Table 11**). The majority of these “rescued” colocalizations were associated with UNC eQTL either not considered significant due to our conservative eQTL calling method or possibly underpowered due to sample size. Most of these eQTL had suggestive nominal p-values but they did not pass our FDR < 0.05 threshold for significance (2.34e-10 ≤ UNC eQTL nominal p-value ≤ 1.17e-3). Nonetheless, these eQTL contained signals robust enough to be considered significant by *coloc*.

### *ABO* and *TNFRSF14* are examples of novel target genes identified using IBD colon tissue

One significant novel colocalization uncovered in colon using IBD tissue was at the 9q34.2 locus associated with CD, recently identified by Liu et al^5^. Our colocalization analysis identified *ABO* (alpha 1-3-N-acetylgalactosaminyltransferase and alpha 1-3-galactosyltransferase) as a potential target gene for this locus (GWAS-UNC PP4 = 0.999). This signal appears to be distinct from non-IBD eQTL for *ABO*, as these eQTL did not colocalize with either GWAS or our UNC eQTL (GWAS-GTEx PP3 = 0.999; GWAS-BarcUVa PP3 = 0.973; UNC-GTEx PP3 = 0.999; UNC-BarcUVa PP3 = 0.518). This signal appeared to be completely absent from BarcUVa, and while this signal was present in GTEx, it did not appear to be the primary driver of *ABO* expression in non-IBD colon (**Figure 3a**). Furthermore, the GTEx online portal reports that their transverse colon data is not predicted to have an eQTL effect with the GWAS index variant in their cross-tissue meta-analysis (METASOFT posterior probability m-value = 0.00).

**Figure 3.**
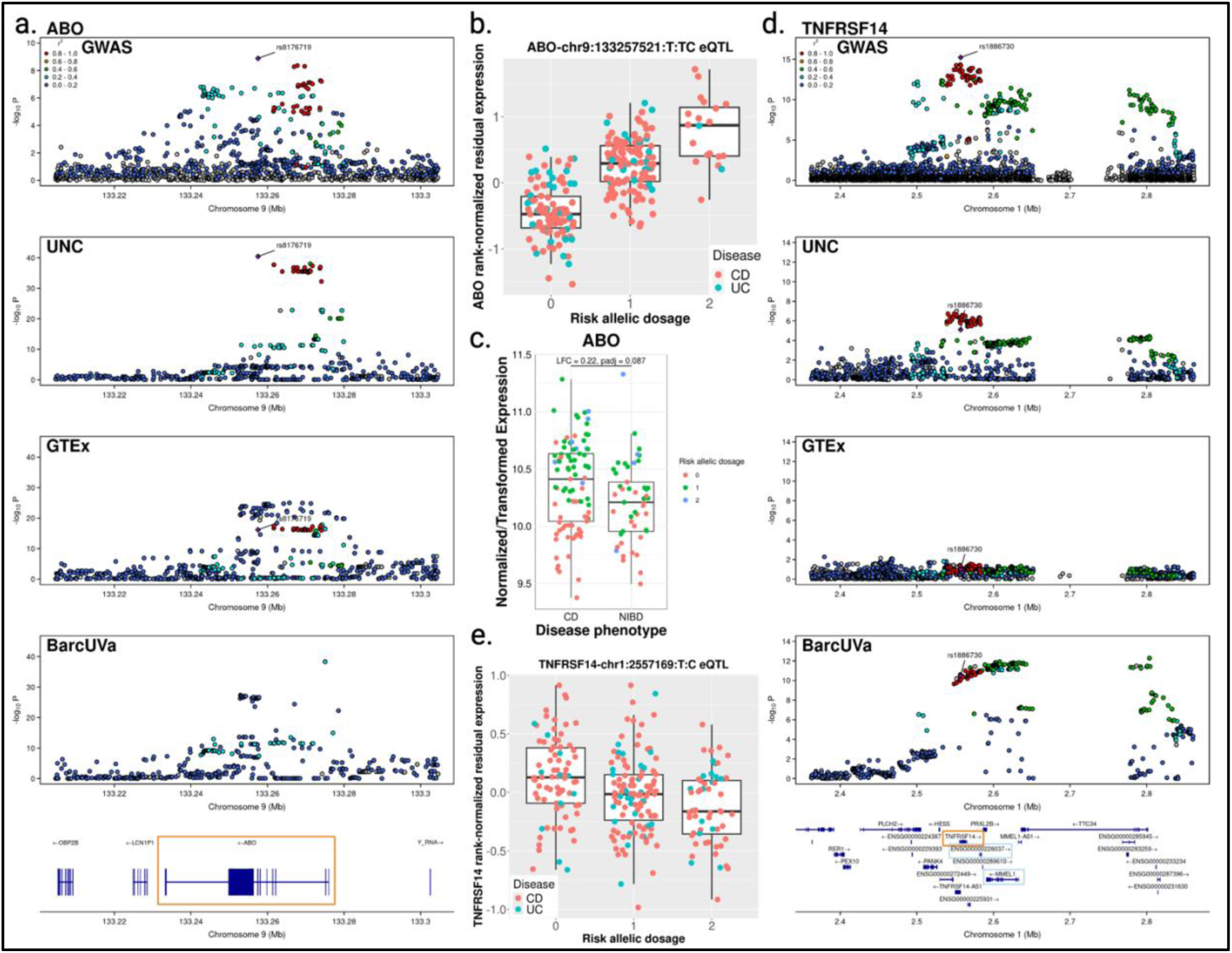
Novel colocalizations and unique target genes are uncovered using diseased tissue. a. Stacked LocusZoom plot for ABO eQTL-GWAS colocalization show shared genetic signals with IBD tissue but not non-IBD tissue. Target gene outlined with orange box. b. ABO UNC eQTL for rs8176719. Each point represents an individual, grouped by risk allelic dosage. Points are colored based on patient diagnosis. c. Boxplot of ABO expression in CD vs non-IBD (NIBD) colon tissue shows trend towards increased expression in CD. Points are colored based on risk allelic dosage for rs8176719. d. Stacked LocusZoom plot for TNFRSF14 eQTL-GWAS colocalization show shared genetic signals using IBD tissue but not non-IBD tissue. Target gene found in diseased tissue outlined with orange box. Other target genes identified using non-IBD tissue outlined in light blue. e. Boxplot of TNFRSF14 UNC eQTL for GWAS index variant rs1886730.

Both our eQTL mapping and GWAS independently identified rs8176719 as the index variant (**Figure 3a**). Rs8176719 is a well-characterized frameshift indel, where the deletion of the C nucleotide leads to a frameshift mutation in exon 6 resulting in early termination^49–51^. This variant is one of the main genetic determinant of histo-blood group antigens, and individuals homozygous for the deletion are blood group O, which has been found to be protective against CD^5^ (**Figure 3b**). We also found that there was a trend towards higher *ABO* expression in CD tissue compared to non-IBD controls (log2 fold change = 0.22, padj = 0.087; **Figure 3c**). In addition to circulating cells, histo-blood group antigens are secreted from and expressed on the mucosal epithelial of the digestive tract, including the colon^52–54^. Secretor status is determined by the *FUT2* locus and has also been linked to CD susceptibilty^30^. These antigens can interact with commensal microbiota and may also play a role in host immunity^24,25^. In fact, this locus has also been linked to gut microbiome composition^55–57^. Liu et al. and others have suggested that blood type may be a potential risk factor for CD, and our colocalization results support this hypothesis by providing the first genetic link to *ABO* expression in the colon^5,58,59^.

In addition, for some GWAS loci, we found colocalizing eQTL from IBD and non-IBD tissue targeted different genes. For example, eQTL from all three data sets colocalized with the 1p36.32 UC risk locus, but the target genes identified by the GTEx and BarcUVa studies were lncRNA *RP3-395M20.7* (GWAS-GTEx PP4 = 0.958) and transmembrane protein-coding gene membrane metallo-endopeptidase like 1 (*MMEL1*; GWAS-BarcUVa PP4 = 0.922), respectively. No known functional evidence supports a link between *MMEL1* and UC, and only limited evidence in a transformed lymphoblastoid cell line suggests *RP3-395M20.7* may play an indirect role^60^.

In contrast, the colocalizing UNC eQTL for this locus was associated with the *TNFRSF14* gene that encodes for a TNF superfamily receptor, also known as the herpesvirus entry mediator (*HVEM*; GWAS-UNC PP4 = 0.914; **Figure 3d**). Interestingly, both GTEx and BarcUVa identified an eQTL for *TNFRSF14*, but in neither case did the eQTL colocalize with GWAS, nor with our UNC eQTL (GWAS-GTEx PP2 = 0.896; GWAS-BarcUVa PP3 = 0.578; UNC-GTEx PP3 = 0.529; UNC-BarcUVa PP3 = 0.520). Like in the *ABO* case described above, the genetic signal seen in the GWAS locus was present in BarcUVa but was not found to be the main regulator of *TNFRSF14* expression in non-IBD colon tissue. Meanwhile in GTEx colon, *TNFRSF14* expression appeared to be regulated by a different signal entirely.

The TNFRSF14/HVEM receptor is unique in that it recognizes multiple distinct ligands including TNFSF14/LIGHT, B and T lymphocyte attenuator (BTLA), lymphotoxin-alpha (LT-α), and CD160, defining an important mediator for immune response homeostasis in the colon^61^. Our colocalization results suggest that decreased *TNFRSF14* is associated with increased risk of UC (**Figure 3e**). While we did not observe a relative difference in colonic *TNFRSF14* expression in either UC or CD patients compared to non-IBD controls, others have found that the absence of Tnfrsf14 accelerated intestinal inflammation in an induced colitis model by transfer of CD4+CD45RB^high^ T cells to *Hvem^-/-^Rag^-/-^* mice^27,28^. Intestinal epithelial Tnfrsf14 signaling has been found to induce epithelial Stat3 activation via NF-κB-inducing kinase, and *Hvem-/-* mice have increased colonic epithelial permeability after intestinal *C. rodentium* infection, suggesting that the expression level of *TNFRSF14/HVEM* is important in epithelial responses and host defense mechanisms in the colon^27^.

### Novel IBD-associated eQTL from diseased tissue have distinct genomic and functional characteristics compared to non-diseased tissue eQTL

Next, we used a variety of approaches to determine whether there were any defining gene regulatory or functional characteristics of the novel and uniquely colocalizing UNC eQTL. In all three studies, uniquely colocalizing eQTL tended to be more distal to the eGene TSS (median absolute distance: BarcUVa eQTL = 64.67 kb; GTEx eQTL = 30.19 kb; UNC eQTL = 41.45 kb) compared to colocalizing eQTL found in all three cohorts (median absolute distance = 14.13 kb). This is consistent with other observations that shared or common eQTL are enriched near promoter regions compared to enhancer-enriched context-specific eQTL^17,62,63^. Interestingly, this distal shift in predicted functional, disease-associated variants was only significant when considering UNC eQTL but not for non-IBD eQTL (Mann-Whitney *U* = 123, p = 0.0114; **Figure 4a**).

**Figure 4.**
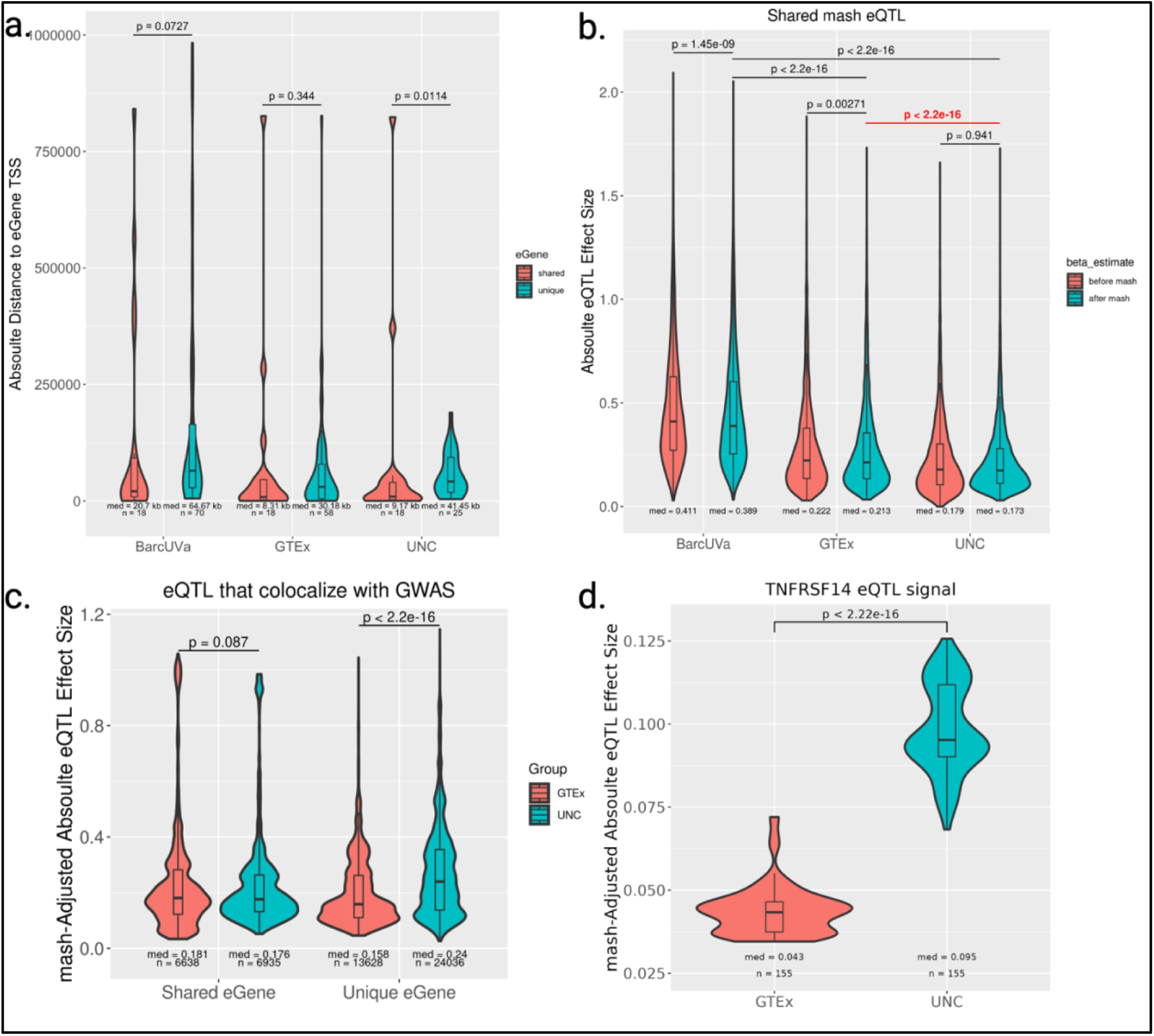
Novel IBD-associated eGenes discovered using diseased tissue have distinct characteristics. a. Violin plots of absolute distance of colocalizing lead eVariants to eGene TSS. Plots are separated by shared eGenes found to colocalize in all three cohorts vs. found uniquely in only one cohort. p-values calculated from Mann-Whitney’s U statistic. b. Violin plots of absolute eQTL effect size distributions for the 11,661 overlapping top eQTL before and after applying mash show UNC eQTL have significantly smaller adjusted effect sizes after mash. p-values calculated from the Mann-Whitney U statistic. c. Violin plots of absolute eQTL effect size distributions for UNC and GTEx eQTL that colocalize with IBD GWAS after applying mash. p-values calculated from Mann-Whitney’s U statistic. d. Violin plot of the mash-adjusted absolute eQTL effect size distribution for overlapping TNFRSF14 eVariants with lfsr < 0.05 show a significantly stronger effect for the UNC eQTL. p-values calculated from Mann-Whitney’s U statistic.

The magnitude of an eQTL effect size represents the relative regulatory difference between alleles, with larger effect sizes representing a greater impact on gene expression. To investigate further how eQTL effect sizes varied across cohorts, we used the “multivariate adaptive shrinkage” (mash) approach. We started by calculating posterior summary statistics for 17,030 eQTL associated with 11,422 eGenes, each representing the top eQTL in at least one study and requiring that they were tested in all three. Of these, mash considered 16,341 eQTL (10,869 eGenes) significant in at least one study using a local false sign rate (lfsr) cutoff < 0.05 (**Supplementary Table 12**). The majority of mash eQTL overlapped across studies and mash effectively shrunk effect size estimates towards zero when appropriate, which increased the pairwise correlation of effect sizes between studies (**Supplementary Figure 8a, 10b**). Importantly, UNC eQTL had the smallest adjusted median absolute effect size of the three data sets, suggesting mash was able to correctly adjust effect sizes that may be inflated due to smaller sample sizes (**Figure 4b**). However, the distributions of absolute mash-adjusted effect size estimates still significantly differed between studies. BarcUVa eQTL had larger estimates than either GTEx or our UNC eQTL, which could help explain the lower levels of sharing reported by mash. This study-specific effect was observed genome-wide, and thus mash preserved this cohort-specific effect (**Supplementary Figure 8c**). Therefore, we only compared effect size estimates between UNC and GTEx eQTL for the remaining analyses.

We compared the mash-adjusted effect sizes for eQTL that colocalized with GWAS and further separated eQTL based on whether the target eGene was commonly found to colocalize in all three cohorts or found to colocalize in only one (i.e. shared or unique; **Supplementary Table 13**). As expected, the absolute effect size distributions for eQTL associated with shared colocalizing eGenes did not significantly differ between UNC and GTEx eQTL. Interestingly, when we considered unique eGenes, the distribution was considerably shifted towards higher adjusted effect sizes for eQTL in diseased tissue (Mann-Whitney *U* = 117337189, p < 2.2e-16; **Figure 4c**). Notably, the direction of this difference in magnitude was reversed compared to the more general set of top eQTL we initially compared, with larger effect sizes in GTEx compared to UNC eQTL (**Figure 4b**). And while we also observed that shared eVariant-eGene pairs between UNC and GTEx eQTL were also significantly shifted, this difference was modest compared to the difference we observed between the uniquely colocalizing effect size distributions (**Supplementary Figure 8d**). Upon closer examination of overlapping variants within individual signals for the 101 shared or unique eGenes, we found 85 of these eGenes had statistically different mash-adjusted effect size distributions between UNC and GTEx eQTL (non-parametric Mann-Whitney U test; **Supplementary Table 14**). Of these, 16 eGenes had consistently larger adjusted effect sizes using diseased tissue, and the majority of which were only discovered using IBD tissue. This set included genes with well-established associations with IBD such as MHC class II genes *HLA-DRB1* and *HLA-DQA1*, as well as some of the novel associations our study has uncovered including *ABO* and *TNFRSF14* (**Figure 4d**).

Finally, we examined the GO terms associated with uniquely colocalizing eGenes to determine whether disease state influenced the types of eGenes discovered to be associated with IBD risk. On average, unique UNC eGenes were associated with more GO terms compared to unique GTEx or BarcUVa eGenes (UNC median = 11 GO terms/eGene; GTEx median = 6 GO terms/eGene; BarcUVa median = 4.5 GO terms/eGene). Most strikingly, we observed a greater proportion of UNC eGenes mapping to the immune response cluster compared to non-IBD eGenes (**Figure 5a-c**). Overall, for both GTEx- and BarcUVa-unique eGenes, the most eGenes fell within the transcriptional regulation by RNA polymerase II (RNAP2) primary cluster with the majority of eGenes mapping to the positive regulation of transcription by RNAP2 secondary cluster in both cohorts. Meanwhile, the primary clusters for transcriptional regulation, cell proliferation, and signal transduction primary clusters equally contained the most IBD-unique eGenes. The signal transduction secondary cluster for G protein-coupled receptor signaling pathway contained the most IBD-unique eGenes, which included multiple genes associated with immune response (*CTSS, HLA-DRB1, IFNGR2, PARK7, TNFRSF14*), and *FERMT1,* which is involved in integrin signaling, an important mechanism for maintaining intestinal homeostasis^64,65^.

**Figure 5.**
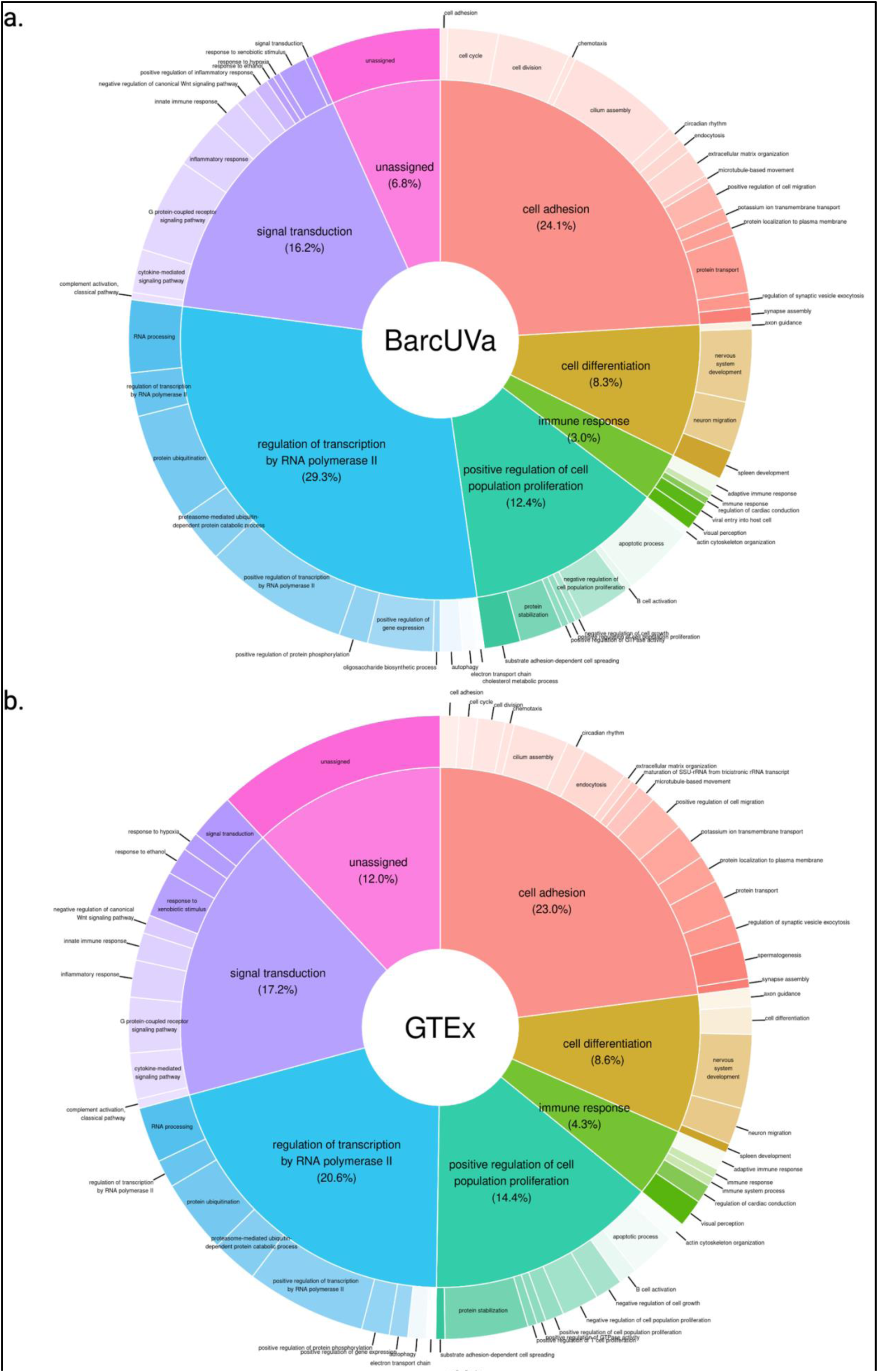
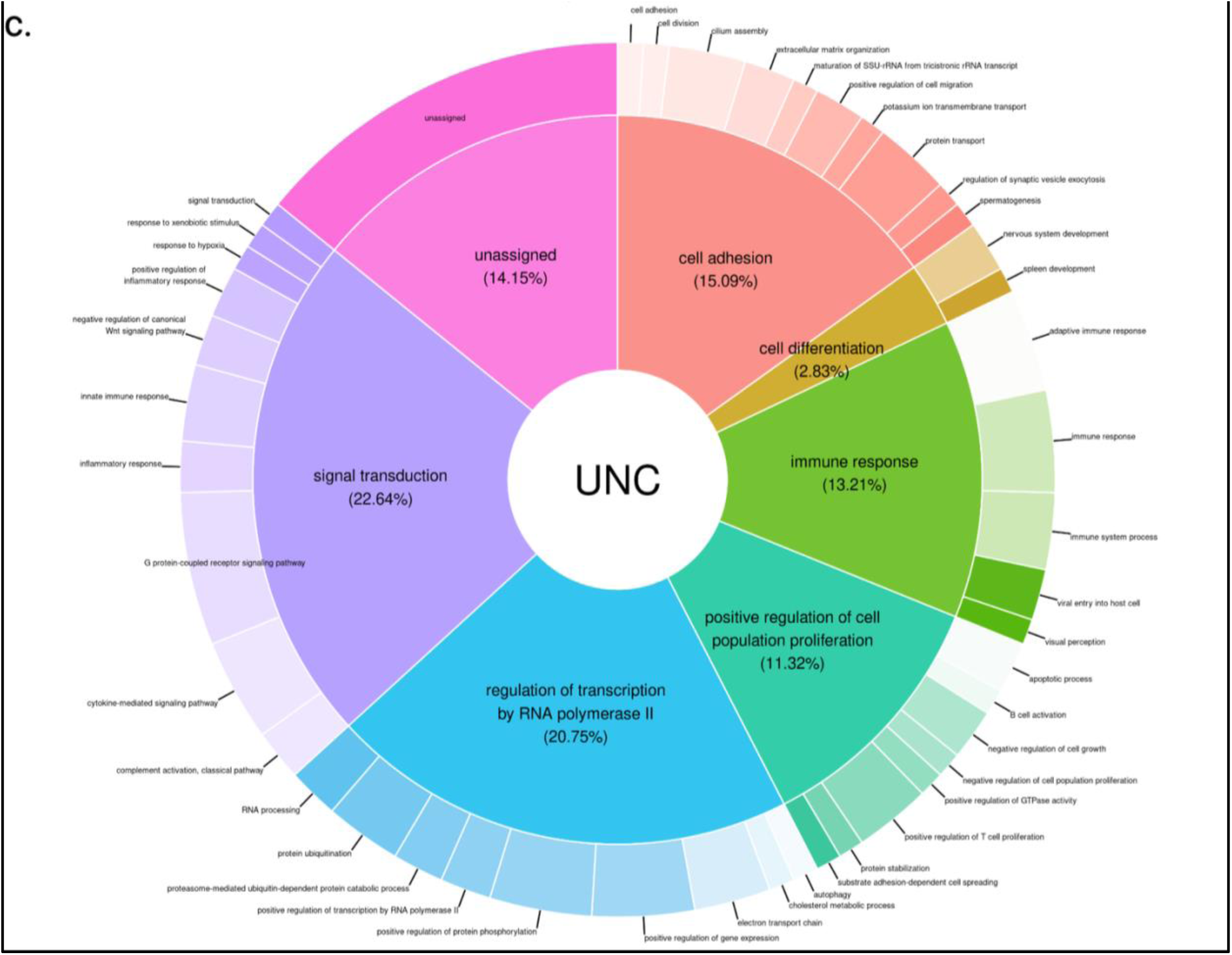
Novel IBD-associated eGenes discovered using diseased tissue are involved in different types of biological processes. a, b, c. Pie-Donut charts representing clustered GO terms associated with uniquely colocalizing eGenes show an increase in the proportion of genes involved with immune response in diseased tissue. Inner pie slices show the proportion of uniquely colocalizing eGenes that map to primary clusters. Outer slices represent proportion of uniquely colocalizing eGenes that map to secondary sub-clusters.

Altogether, these results provide important insights into the unique regulatory characteristics of eQTL in diseased tissue, particularly in the context of IBD. Furthermore, these findings highlight how disease context can reveal regulatory mechanisms and effects, like immune responses, that may remain hidden or be challenging to detect without environmental triggers that ultimately lead to disease.

## Discussion

Here we report the most comprehensive set of 194 potential target genes for 108 IBD-associated loci using colon eQTL from both IBD and non-IBD individuals. We found that these target genes are associated with a diverse range of biological functions relevant to IBD pathogenesis including cell adhesion, differentiation, proliferation, immune response, transcriptional regulation, and signal transduction. They include both genes previously related to IBD for which we now have evidence of genetic influence, as well as genes that have not been explored in connection to changes in the intestinal environment known to contribute to IBD.

Importantly, we have shown that eQTL mapped using tissue from patients with IBD were more strongly enriched for IBD-associated variants compared to non-IBD tissue and included a unique subset of eQTL that were not found in current non-IBD colon eQTL data sets. This has also allowed us to identify additional target genes for IBD GWAS loci compared to only focusing on non-IBD cohorts. For example, previous research has suggested an association between ABO blood groups with IBD, but we are the first to show that genetic variation may also modulate *ABO* expression in the colon, contributing to its role in disease^5,58,59^. We also identified *TNFRSF14* as a target gene in an IBD locus in which other nearby genes were prioritized in non-IBD eQTL data. Prior studies have clearly shown that reduced TNFRSF14 expression contributes to intestinal inflammation and more severe IBD-related phenotypes, but evidence for a genetic linkage to its regulation in colon has not been previously reported^26–28^. Findings such as these suggest that leveraging diseased tissue can provide more accurate hypotheses for functional validation, saving both time and money.

These results also suggest that the disease state can alter the regulatory landscape that explain GWAS loci. The larger effect sizes for novel IBD-associated eQTL in diseased tissue suggest that disease state may amplify the genetic effects on gene expression which are less apparent in non-diseased tissue. This is underscored by potential stronger biological impact associated with the types of target genes identified using IBD tissue. This contrasts with the model recently proposed by Mostafavi et al., who suggest that disease-critical eQTL are rare in normal tissue and characterized by small changes in gene expression tolerated by natural selection^66^. However, we identified several of the genomic and functional characteristics in our novel eQTL that align with their characterization of GWAS genes. By actively focusing on diseased tissue, increased effect sizes of certain disease-relevant eQTL will facilitate their discovery and support their relevance to disease without necessitating the need for impractically large sample sizes. With increased IBD patient sample sizes and exploration of other relevant tissues, we expect to detect additional and potentially previously undetected associations.

Bulk tissue-based eQTL represent genetically regulated signals averaged across multiple cell types. The colonic mucosa includes multiple epithelial and immune cell types that play direct roles in regulating the intestinal barrier and immune responses associated with IBD. Our eQTL model, though, allowed us to report gene regulatory effects without dependency on cell type abundances. However, we acknowledge that our study cannot identify specific cell types in which target gene expression or particular pathways may have the greatest impact on disease risk. eQTL studies using single-cell data are currently underpowered, making a combination of bulk tissue and single-cell studies the most viable solution for now. Comparing and combining independent eQTL studies is also challenging due to limited access to individual-level data and reliance on summary data. We performed multiple parallel statistical comparisons to support our conclusions. We acknowledge that to more definitively determine whether disease state alters regulatory mechanisms for some genes resulting in new eQTL or increased eQTL effect sizes, we need better powered studies in both diseased and non-diseased tissue, ideally within the same study.

eQTL studies, such as the one reported here, provide strong evidence for target genes within GWAS loci. While this is an important step forward, eQTL are unable to conclusively identify casual variants and gene regulatory mechanisms. Linking other molecular measures such as chromatin accessibility and other epigenetic markers to genetic variation will be able to address this question more directly. A concerted effort in conducting these studies in tissues from both affected and unaffected individuals will be important to disentangle the regulatory landscape and the role of common genetic factors contributing to IBD.

## Methods

### Patient recruitment and sample collection

This study was approved by Human Research Ethics Committee at UNC-Chapel Hill and carried out with accordance to 10-0355 and 11-0359. Patients were recruited at the UNC Multidisciplinary Center for IBD. Written informed consent was obtained from all participants. Approximately 52% of patients included in this study were female and the median age at sample collection was 38 ± 14.98 years. Non-inflamed mucosal samples were collected from primarily ascending colonic tissue from IBD patients undergoing colonoscopy or surgical resection. Tissue samples were immediately submerged in RNAlater and snap frozen in liquid nitrogen to prevent RNA degradation. Peripheral blood samples were also collected for DNA analysis. Additionally, we included data from 61 CD patients that reside in the Crohn’s & Colitis Foundation IBD Plexus platform and were collected from the 20cm mark above the rectum as part of SPARC IBD. Patients were consented and biosamples were collected and processed as previously described^67^. Samples from a total of 252 IBD patients (CD n = 199; UC n = 53) were selected for downstream analyses after genotype and RNA-seq QC.

### Genotyping and Imputation

Genomic DNA was extracted from whole blood or tissue biopsies for genotyping array. DNA was genotyped for > 650,000 markers using the Infinium Global Screening Array-24 BeadChip (v3.0 or v1.0, Illumina). Markers were mapped to hg38 or lifted over from hg19 using *CrossMap* (v0.6.3, Python v3.6.6). We performed quality control on genotyped samples using plink (v1.90b3). Samples with a missingness rate >10% of genotype calls or within two degrees of relatedness were removed. Additional genotypes were imputed using TOPMed Imputation Server with the TOPMed-r2 panel and phasing was performed using Eagle (v2.4). Sites with an imputation quality of R^2^ < 0.3 were discarded, resulting in a total of 27,569,639 SNPs and 2,075,971 indels. Variants were annotated with rsIDs from dbSNP build 156.

#### Genetic similarity inference

To control for population structure effects, principal component analysis (PCA) was run on genotype data using R (v4.1.3). We restricted the analysis to a selection of LD-pruned (R^2^ < 0.2) typed autosomal sites meeting the following criteria using plink (v1.90b3): genotype missingness < 0.05 and MAF > 0.05. Genotypes were then compared to 1000 Genomes reference samples to infer genetic similarity of IBD patients to 1000 Genomes super populations. We identified 84.52% of patients used in this study to have the greatest genetic similarity to the 1000 Genomes reference European superpopulation and 13.10% of patients to have the greatest genetic similarity to the African superpopulation (**Supplementary Figure 9**).

### RNA sequencing and gene quantification

Total RNA was extracted from primarily colonic mucosa samples for paired-end RNA-sequencing with an average sequencing depth of 41.8M reads. Reads were aligned to the human genome (hg38/GRCh38) based on GENCODE v39 annotations using STAR (v2.7.9a). To prevent allelic mapping biases, reads were aligned using the “--WaspOutputMod” with corresponding sample genotypes when running STAR. We used *verifyBAMID* (v1.1.3) to ensure proper pairing between RNA-sequencing and genotype samples. Gene expression was quantified using QTLtools (v1.3.1).

### Removing unwanted variation (RUV) from expression data

We applied the RUVSeq method to remove unwanted variation in normalized RNA-seq expression data and to account for hidden technical and biological confounders as an alternative to using the probabilistic estimation of expression residuals (PEER)^68,69^. RUVSeq uses factor analysis to adjust for unwanted variation applied to count data from negative control genes (i.e. genes not expected to be influenced by the biological factor of interest). Briefly, raw count data were filtered to remove genes with low coverage, requiring at least 5 raw reads in at least 25% of samples. Counts were normalized and transformed using DESeq2’s vst() function for variance stabilizing transformation^70^. Known covariates batch, sex, and the first four genotype PCs were regressed from the normalized count data using limma’s removeBatchEffect(). Using the log-scale normalized, transformed, and covariate-corrected counts, we selected the top 1000 genes with the smallest coefficient of variation out of the top 5000 most highly expressed genes as our negative control gene set for RUV factor estimation. The initial number of RUV factors calculated was determined by the total sample size, *n*, where the number of estimated RUV factors is *k* = 0.25*n*, as recommended by developers of PEER.

The number of RUV factors included in the final eQTL model was determined using the kneedle algorithm from the *kneed* package (v0.5.0; Python v3.6.6)^71^. Kneedle calculates the point of maximum curvature by finding the local maxima of the difference curve from *y = x* after smoothing and normalizing the data. This is a conservative quantitative method similar to finding the elbow or “knee” of the curve that maintains the overall behavior of the curve and limits false positives. eQTL models were iteratively run in increments of +1 RUV factors added to each additional model as a covariate to determine the number of unique genes with significant eQTL (eGenes) detected using a false discovery rate (FDR) of 5%. We plotted the number of significant eGenes by the number of RUV factors included in each model (**Supplementary Figure 10**).

### *cis*-eQTL mapping using IBD tissue

We used genotype and gene expression data from 252 IBD patients to map *cis*-eQTL within ± 1 Mb of the transcription start site (TSS) of each gene using QTLtools (v1.3.1)^72^. We tested variants with a minor allele frequency (MAF) ≥ 0.02 in our patients and gene trimmed means of M values were rank inverse normal transformed. The final linear model was adjusted for sex, batch, population structure using the first four genotype PCs, and 21 RUV factors. Local adjusted p-values were determined using a null beta distribution generated using 1000 permutations per test. Global p-values were adjusted for multiple hypothesis testing correction using the Storey and Tibshirani procedure for FDR^73^. eQTL with an FDR < 5% were considered significant.

### Comparison to non-IBD eQTL studies

Publicly available eQTL summary statistics for GTEx (v8) and BarcUVa colon tissue were downloaded^12,13^. For GTEx, we focused our main analyses on the transverse colon tissue only as the sigmoid colon tissue was collected from the muscularis propria. BarcUVa variant coordinates were lifted over to hg38 using the liftOver function within the *rtracklayer* R package (v1.54.0). For comparison across eQTL data sets, variants were matched based on coordinate position, and reference and alternative alleles and genes were matched based on Ensembl gene IDs excluding version numbers. Pairwise effect size estimates were correlated using Pearson’s correlation coefficient and scatter plots were created using R (v4.1.3). Storey’s π1 estimates were calculated using nominal p-values from GTEx or BarcUVa using the *qvalue* R package (v2.26.0) to estimate the proportion of true positives in our eQTL data set.

#### Colocalizing eQTL signals across studies

We colocalized primary eQTL signals for shared eGenes across colon eQTL studies to detect shared genetic signals using the *coloc* R package (v5.2.3)^74^. We created a union set of genes tested for eQTL in all three studies, and colocalized signals for which there was a significant eQTL present in at least one study. We tested all pairwise comparisons with available summary statistics. eQTL-eQTL pairs with a H4 posterior probability (PP4) > 0.5 were considered colocalized. LocusZoom plots were generated using the *locuszoomr* R package (v0.2.0) and Ensembl annotation v105.

#### eQTL effect size analysis

We applied multivariate adaptive shrinkage (mash) to quantify the sharing of eQTL signal effect size estimates across disease states using the *mashr* R package (v0.2.79)^75^. Mash uses an empirical Bayes approach to estimate the patterns of sharing across conditions and uses these patterns to improve effect size estimates, which can then be used to assess effect size heterogeneity more quantitatively across conditions. We followed the eQTL analysis workflow outlined by the developers (https://stephenslab.github.io/mashr/index.html). Briefly, we aggregated eQTL effect size estimates and standard errors tested in all three colon data sets into matrices with over 53M eVariant-eGene features which were used as input for mash analysis. We selected the most significant eQTL for these eGenes across all conditions as our set of high confidence eQTL, referred to as our “strong” set. We used a set of 5M randomly selected eVariant-eGene pairs tested in all three data sets to calculate the null correlation. We fit the mash model to the random set using a set of data-driven covariances learned from the “strong” set to estimate the mixture proportions. We then used this fit to calculate the posterior summaries for the “strong” set and for eQTL that colocalized with GWAS. We focused our analyses on eQTL with a local false sign rate (lfsr) < 0.05, as recommended. We used the Mann-Whitney *U* test and Pearson’s correlation coefficient calculated in R to compare each eQTL signal separately. P-values were adjusted for multiple hypothesis testing correction using the BH-procedure through the *qvalue* package (v2.26.0).

### Colocalizing eQTL with GWAS

eQTL were colocalized with GWAS loci using a two-stage approach. We first calculated the linkage disequilibrium (LD) between GWAS index variants and primary lead eVariants across all three eQTL data sets using the summary statistics from the recent meta-analysis published by Liu et al. LD proxies for GWAS index variants were queried across all reference populations using the TOPLD API^76^. eQTL-GWAS pairs with an LD R^2^ > 0.2 in at least one reference population were prioritized for colocalization testing using *coloc*. Full summary statistics were used to calculate statistical colocalization of genetic signals using the default priors. eQTL-GWAS pairs with a H4 posterior probability (PP4) > 0.5 were considered colocalized. LocusZoom plots were generated using the methods described above.

To identify novel GWAS colocalizations, we first identified the unique set of colocalizing eGenes within our IBD eQTL data set with PP4 > 0.5. We then employed a “cross-colocalization” approach: the corresponding non-IBD summary statistics for these eGenes were then tested for GWAS colocalization, regardless of either eQTL significance or LD-based prioritization, to determine whether the colocalized signal was also present in non-IBD samples. We considered eQTL with highest posterior probabilities for either H2 (*coloc* detect a signal in GWAS only) or H3 (*coloc* detects signals in both eQTL and GWAS with distinct causal variants) in both GTEx and BarcUVa to be novel within our IBD eQTL data set. eQTL with the highest posterior probability for H4 in at least one non-IBD data set were considered shared.

### GWAS enrichment analysis

We calculated the overrepresentation of eVariants among GWAS variants using an approach similar to one reported by the GTEx Consortium^12^. For each trait (CD, UC, IBD), we extracted all significant GWAS variants using a genome-wide threshold p < 5e-8. Then for each tissue or study, we compared the proportion of significant eVariants (FDR < 5%) among GWAS variants to the proportion among all tested variants. We then calculated the enrichment fold of eVariants among GWAS variants versus the baseline of the proportion of eVariants among all tested variants. A heatmap of enrichment scores was created using the *pheatmap* package in R (v1.0.12).

### Gene ontology (GO) term clustering

GO terms were queried for eGenes associated with GWAS loci through colocalization using the Database for Annotation, Visualization, and Integrated Discovery (DAVID, v2024q1)^77,78^. We built a hierarchy of clustered GO terms associated with colocalizing eGenes using the *rrvgo* package in R(v1.6.0)^79^. Briefly, pairwise semantic similarity scores were calculated using a graph-based method that takes advantage of the topology of the GO graph structure between two GO terms^80^. Hierarchical clustering with complete linkage was used to group GO terms based on similarity scores. We created a three-tier hierarchy of clustered GO terms using a recursive clustering approach, using a threshold = 0.99 for primary term clusters and 0.90 for secondary clusters. These thresholds were used in combination with GO term set sizes to determine clusters and to favor broader biological terms to serve as cluster representatives. Clustered GO terms were then mapped back to the eQTL data sets and visualized using pie-donut charts created using the *webr* package in R (v0.1.5).

### Differential expression analysis

Differential gene expression analysis was conducted using DESeq2 (v1.34.0) in R (v4.1.3) to assess relative fold change differences between IBD and non-IBD colonic gene expression^70^. For controls, we included RNA-seq data generated from non-IBD patients at UNC and sequenced with IBD samples used in above eQTL analyses. Non-IBD samples were processed as outlined above. Raw counts were imported into DESeq2 and transformed using the variance stabilizing transformation. RUV factors were calculated using a set of non-differentially expressed autosomal genes (p > 0.1) and included in the generalized linear model along with sex and batch as covariates to identify differentially expressed genes between IBD and non-IBD samples. We conducted the following comparisons: CD vs. non-IBD and UC vs. non-IBD. Genes with an FDR-adjusted p-value < 0.05 were considered differentially expressed.

### Statistical Analyses

Unless otherwise noted, all statistical tests were performed using R (v4.1.3).

## Data Availability

Normalized gene count data are available in the Gene Expression Omnibus (GEO) accession number GSE279302. Raw RNA-seq and genotypes are available in dbGaP. IBD Plexus data are available upon approved application to Crohn’s & Colitis Foundation IBD Plexus program (https://www.crohnscolitisfoundation.org/ibd-plexus). Publicly available v8 eQTL summary statistics were downloaded from the GTEx portal (https://gtexportal.org/home/downloads/adult-gtex/overview). Full summary statistics for BarcUVa-seq eQTL were downloaded from the Digital Repository of the University of Barcelona (https://diposit.ub.edu/dspace/handle/2445/172697). GWAS summary statistics can be downloaded from https://www.ibdgenetics.org.

## Code Availability

eQTL pipeline is available on GitHub: https://github.com/nnishiyama/IBD_colon_eQTL

## Acknowledgements

The authors thank the participants, clinicians, and investigators of the Study of a Prospective Adult Research Cohort with IBD (SPARC IBD) who have contributed time, data, and samples. SPARC IBD is a part of the Crohn’s & Colitis Foundation’s IBD Plexus data-sharing platform that includes data from electronic health records and biospecimens. This study was supported by the National Institute of Diabetes And Digestive And Kidney Diseases of the National Institutes of Health (NIH) Award Numbers F31DK137574, 2P01DK094779, 1R01DK136262, 1R01DK138462; The Leona M. and Harry B. Helmsley Charitable Trust Award Number 2105-04679; and NIH’s National Institute of General Medical Sciences Award Number T32-GM135123. We gratefully acknowledge the technical support from the UNC High Throughput Sequencing Facility and the Mammalian Genotyping Core. These facilities are supported by the University Cancer Research Fund, Comprehensive Cancer Center Core Support grant (P30-CA016086) and the UNC Center for Mental Health and Susceptibility grant (P30-ES010126).

## Author Contributions

T.S.F. and S.Z.S. conceptualized study. M.R.K, R.R., T.S.S., J.S., J.F., and W.A.B. contributed samples. M.B., C.B., G. Lau, and G. Lian assisted in sample acquisition. N.C.N., S.S., K.D., A.B., K.C., R.G., M.M.K.N., and M.R.S. contributed to data generation and analysis. N.C.N. and T.S.F. provided data interpretation. N.C.N, S.Z.S, and T.S.F. prepared manuscript.

## Supplementary Figures

**Supplementary Figure 1.**
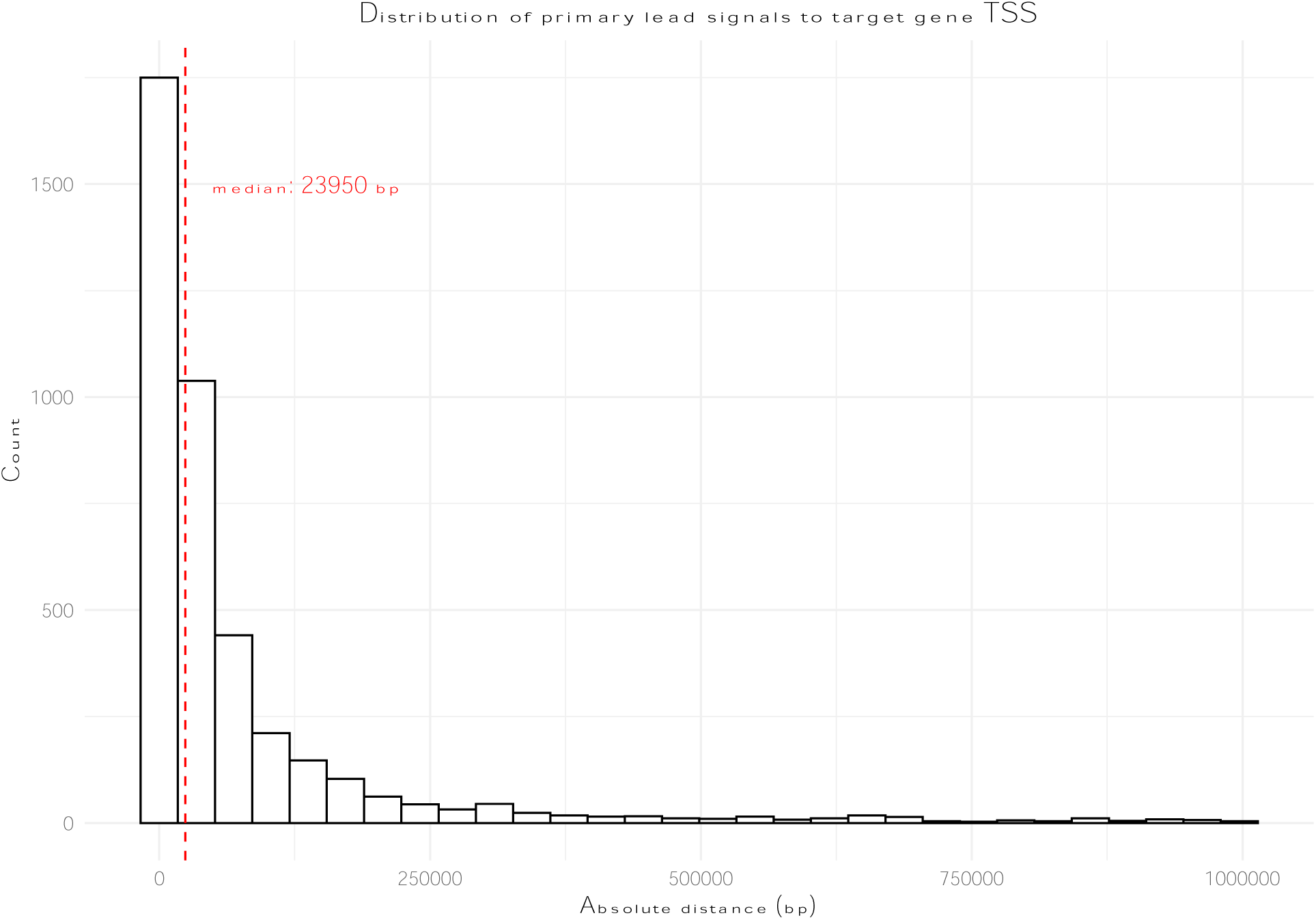
Primary lead eQTL signals are proximal to target gene transcription start site (TSS).

**Supplementary Figure 2.**
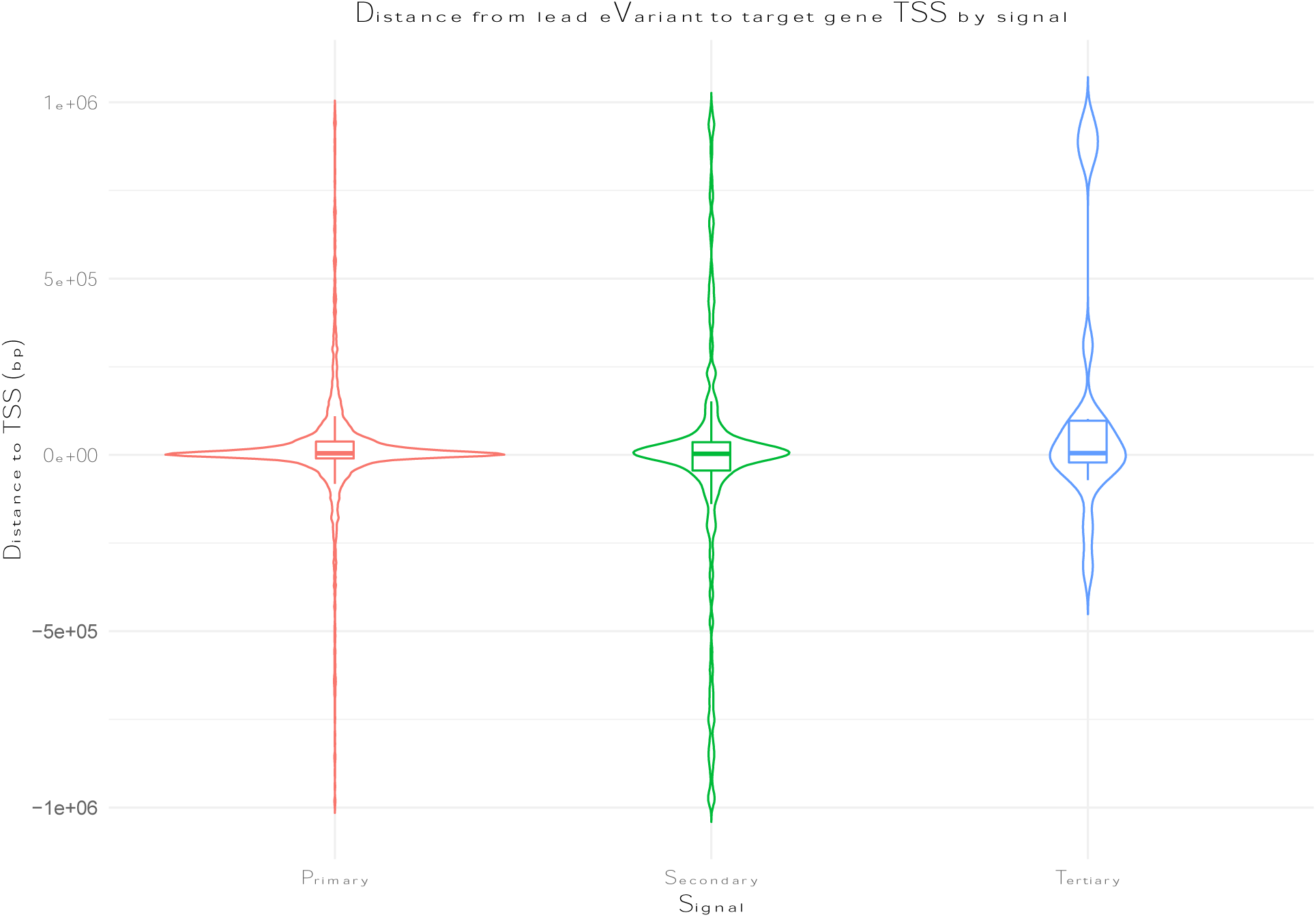
Distribution of distance from lead eVariant to target gene TSS by signal rank.

**Supplementary Figure 3.**
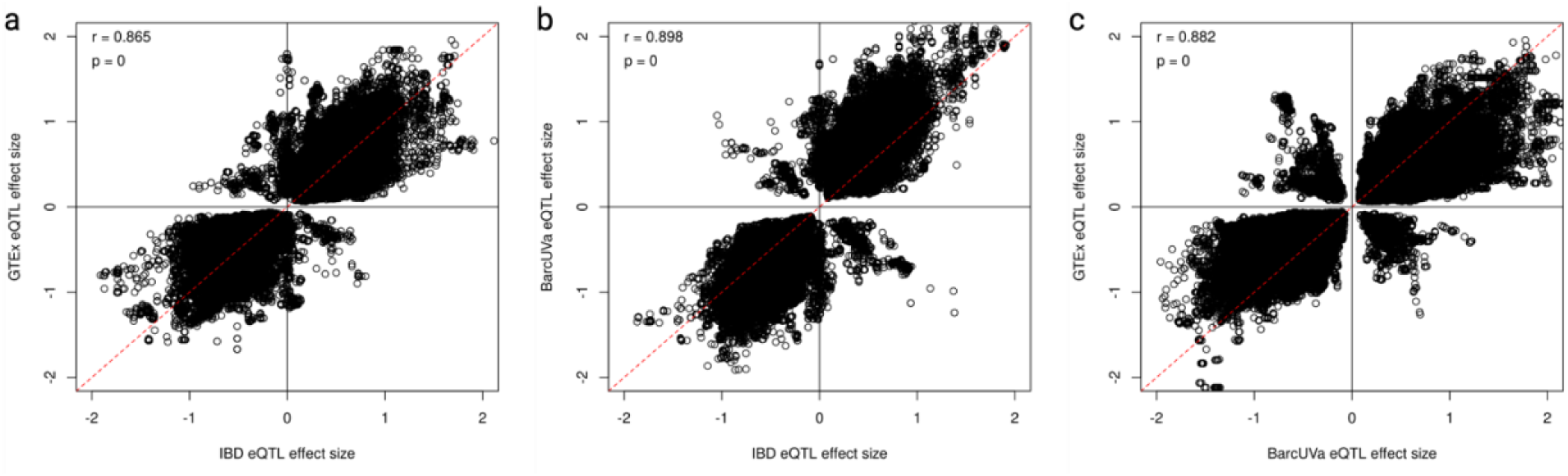
Effect Size Correlation between all significant shared eVariant-eGene pairs.

**Supplementary Figure 4.**
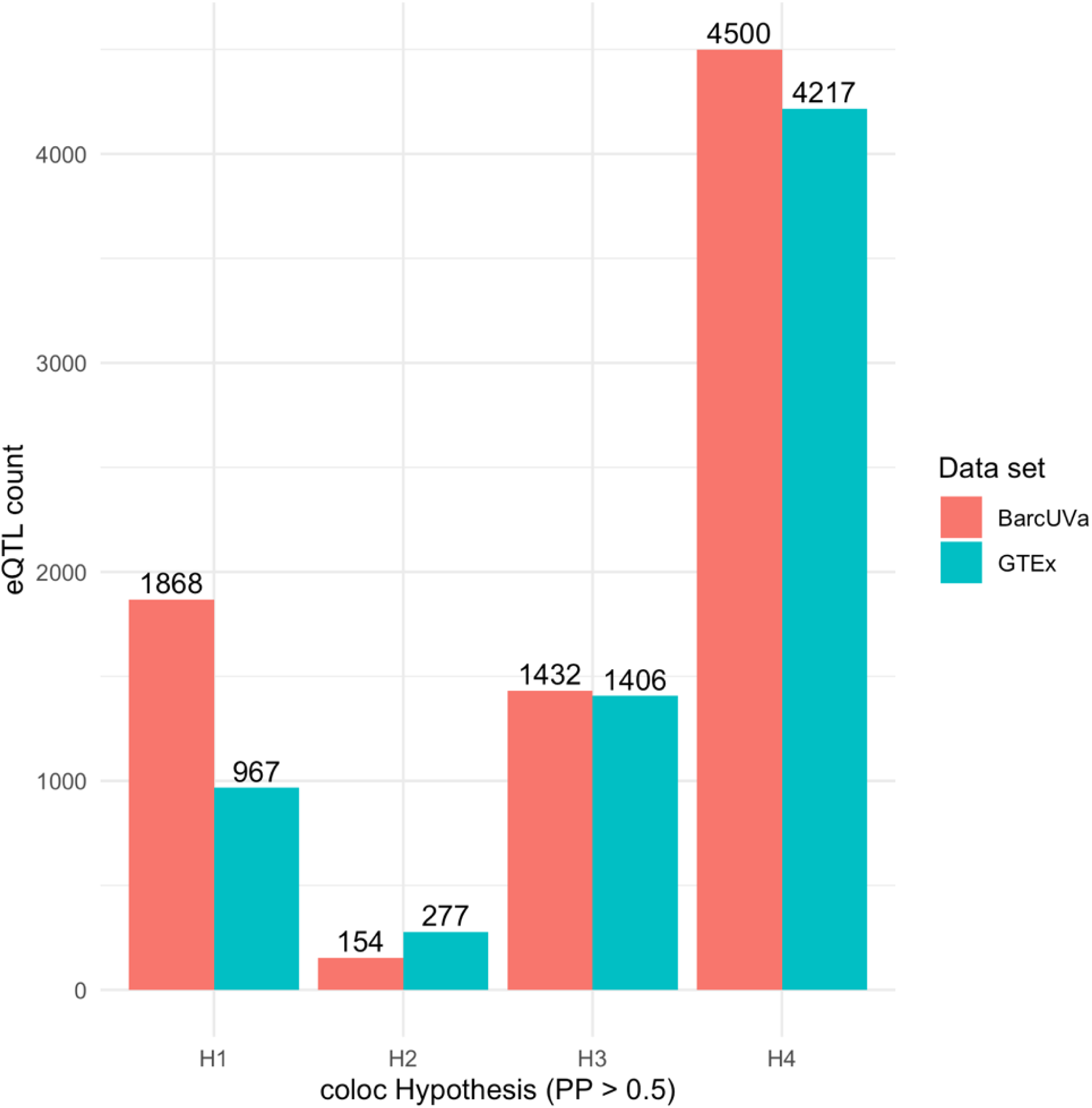
Bar plot of GTEx eQTL-BarcUVa eQTL colocalization results by coloc hypothesis (PP > 0.5): H1, significant signal detected in reference data set only (self); H2, significant signal detected in comparison data set only; H3, significant signal detected in both data sets but different causal variants; H4, significant signal detected in both data sets, same causal variant.

**Supplementary Figure 5.**
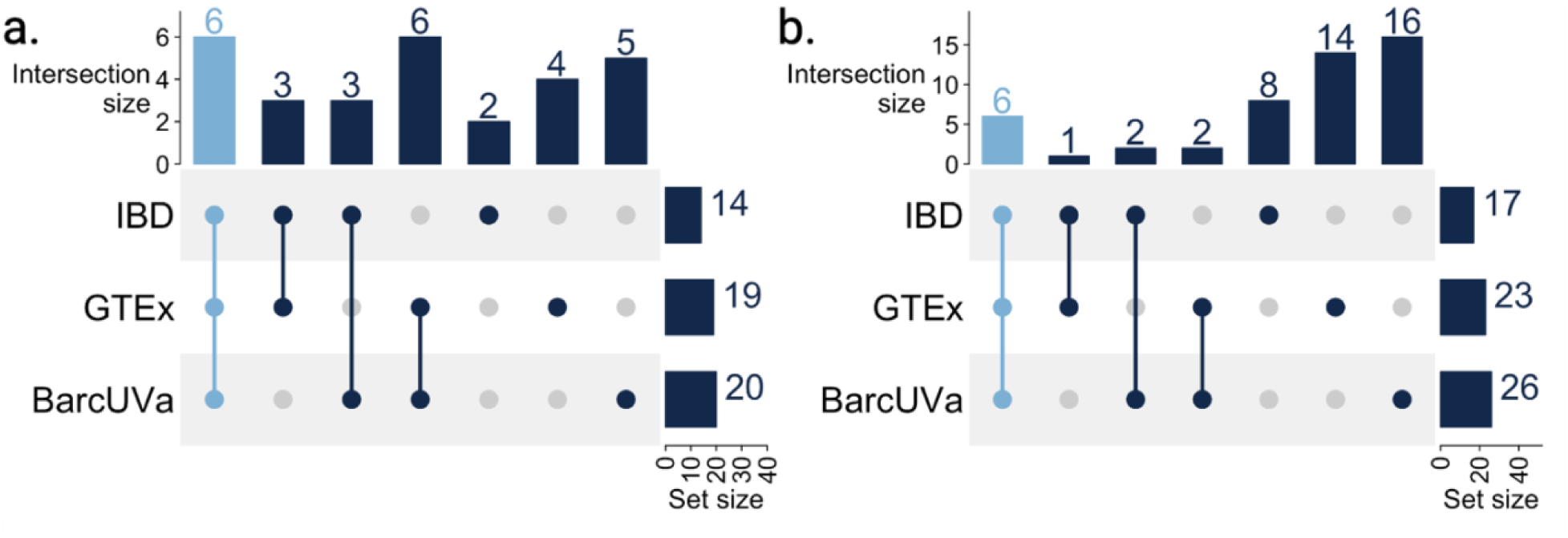
Sharing of colocalization results for newly reported IBD GWAS loci. a. UpSet plot showing overlap of newly reported colocalizing GWAS loci across studies. b. UpSet plot showing overlap of eGenes that colocalize with newly reported IBD GWAS loci across studies.

**Supplementary Figure 6.**
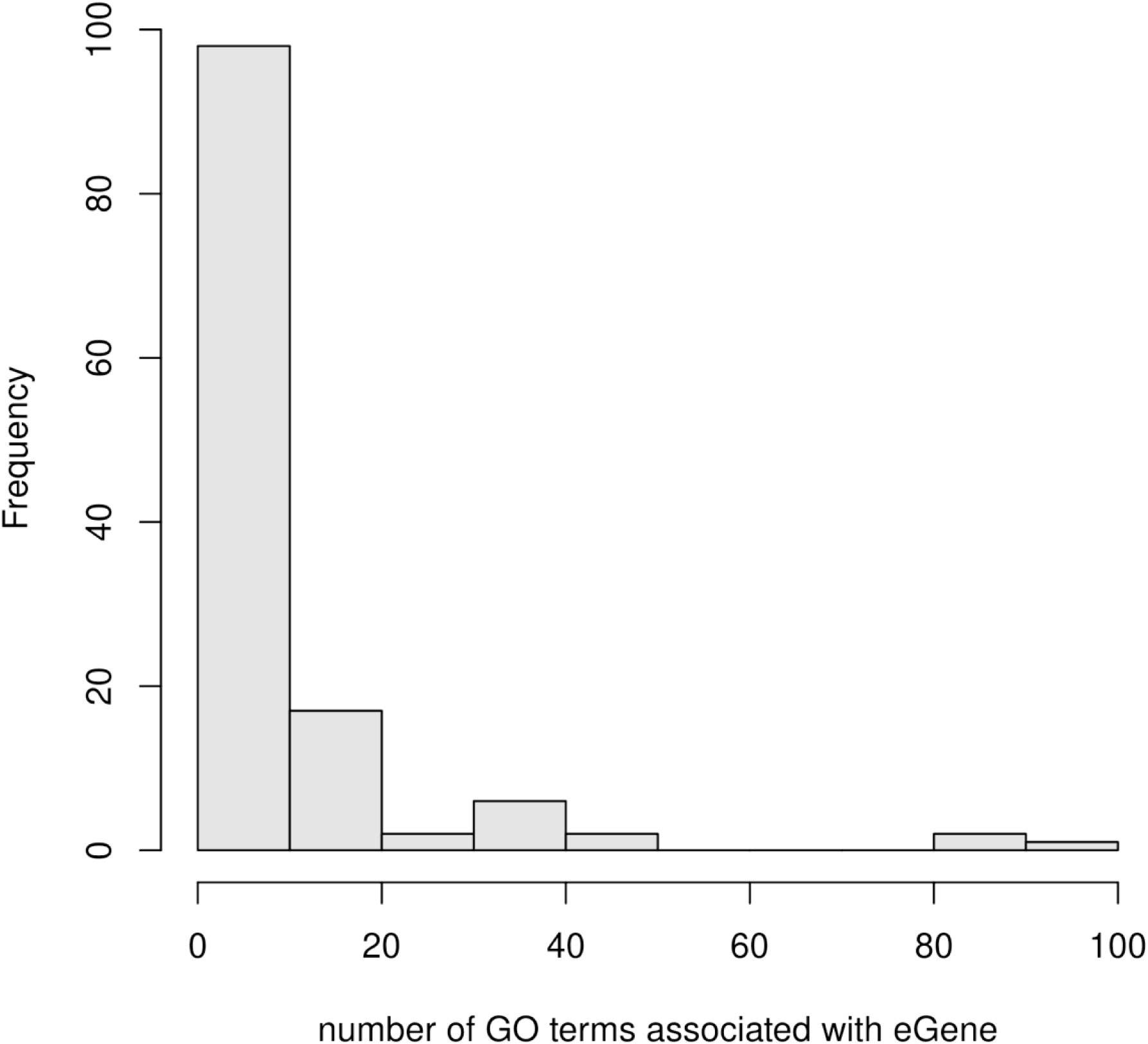
Histogram of the number of GO terms associated with colocalizing eGenes.

**Supplementary Figure 7.**
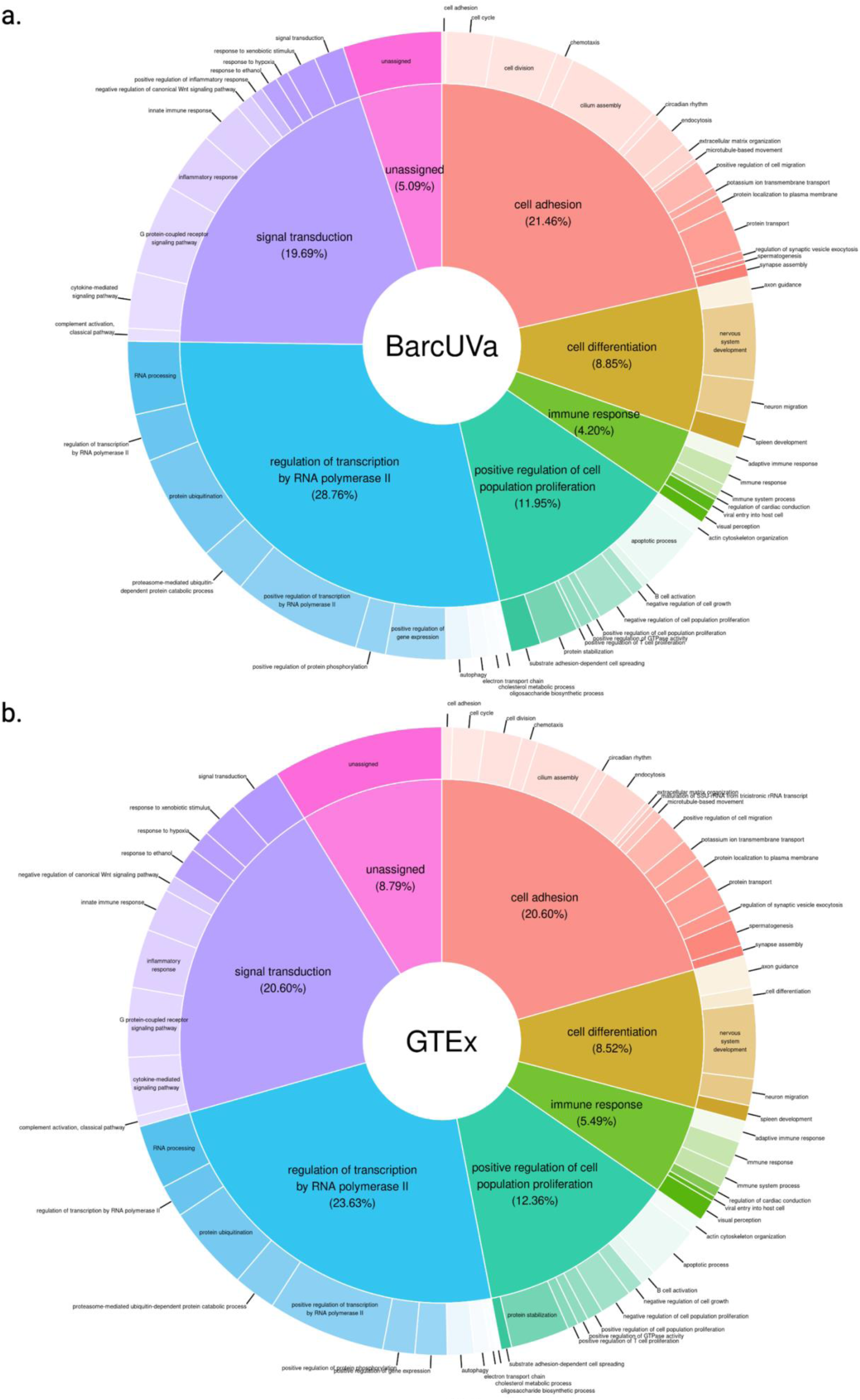
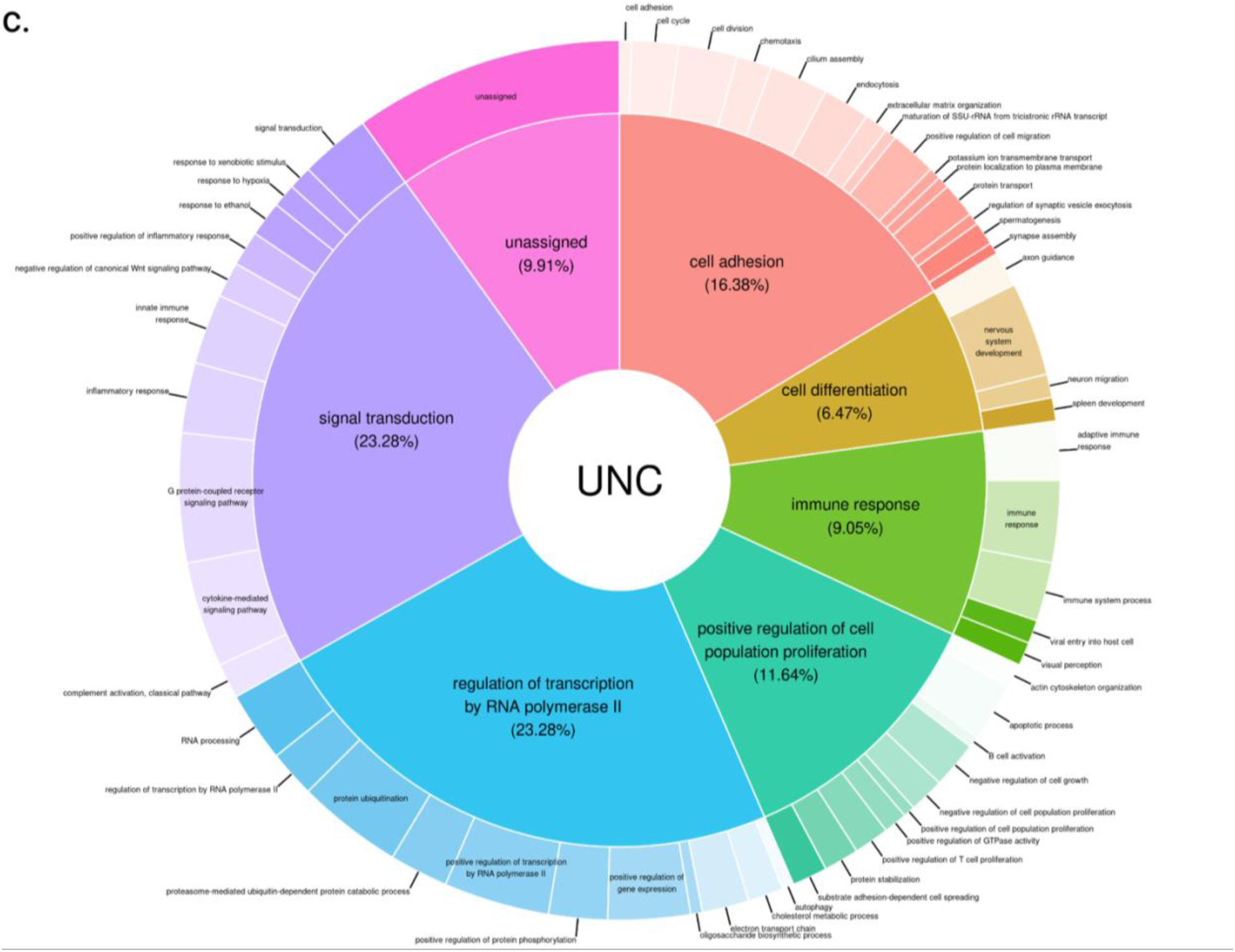
Pie-Donut charts representing clustered GO terms associated with colocalizing eGenes. Inner pie slices show the proportion of colocalizing eGenes that map to primary clusters. Outer slices represent proportion of colocalizing eGenes that map to secondary sub-clusters.

**Supplementary Figure 8.**
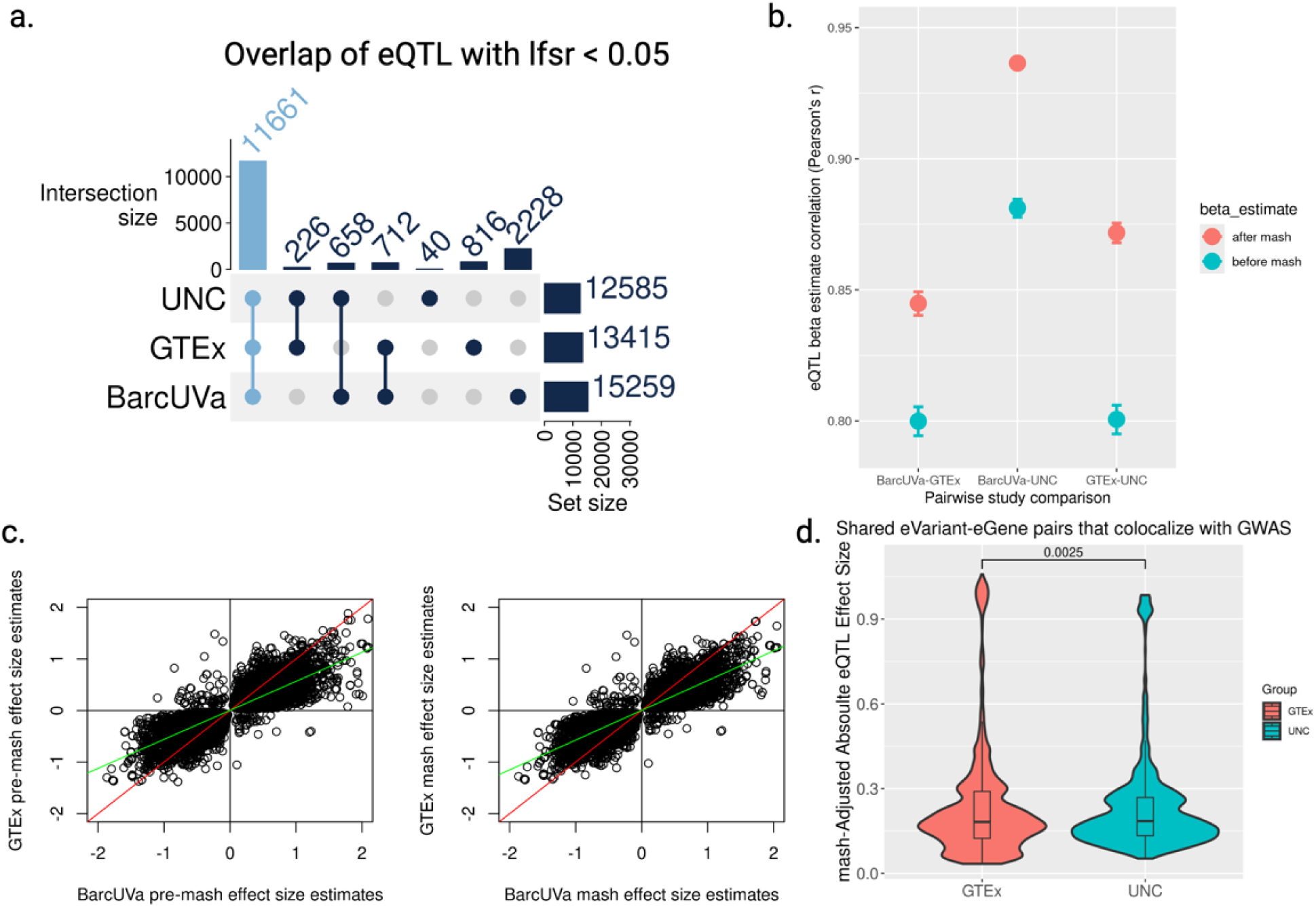
Mash results. a. UpSet plot of overlap for eQTL with lfsr < 0.05. The 11,661 overlapping eQTL highlighted in Carolina blue were used to generate data in panels b and c. b. Scatterplot of correlation estimates for pairwise eQTL study comparisons of effect size estimates before and after applying mash. Vertical bars represent the 95% confidence interval for each estimate. c. Effect size correlation for non-IBD eQTL before and after applying mash. The red line represents the (0,1) intercept. The green line represents the regression fit. d. Violin plots of absolute eQTL effect size distributions for the 6315 shared eVariant-eGene pairs that colocalize with GWAS loci. p-value calculated from the Mann-Whitney U statistic.

**Supplementary Figure 9.**
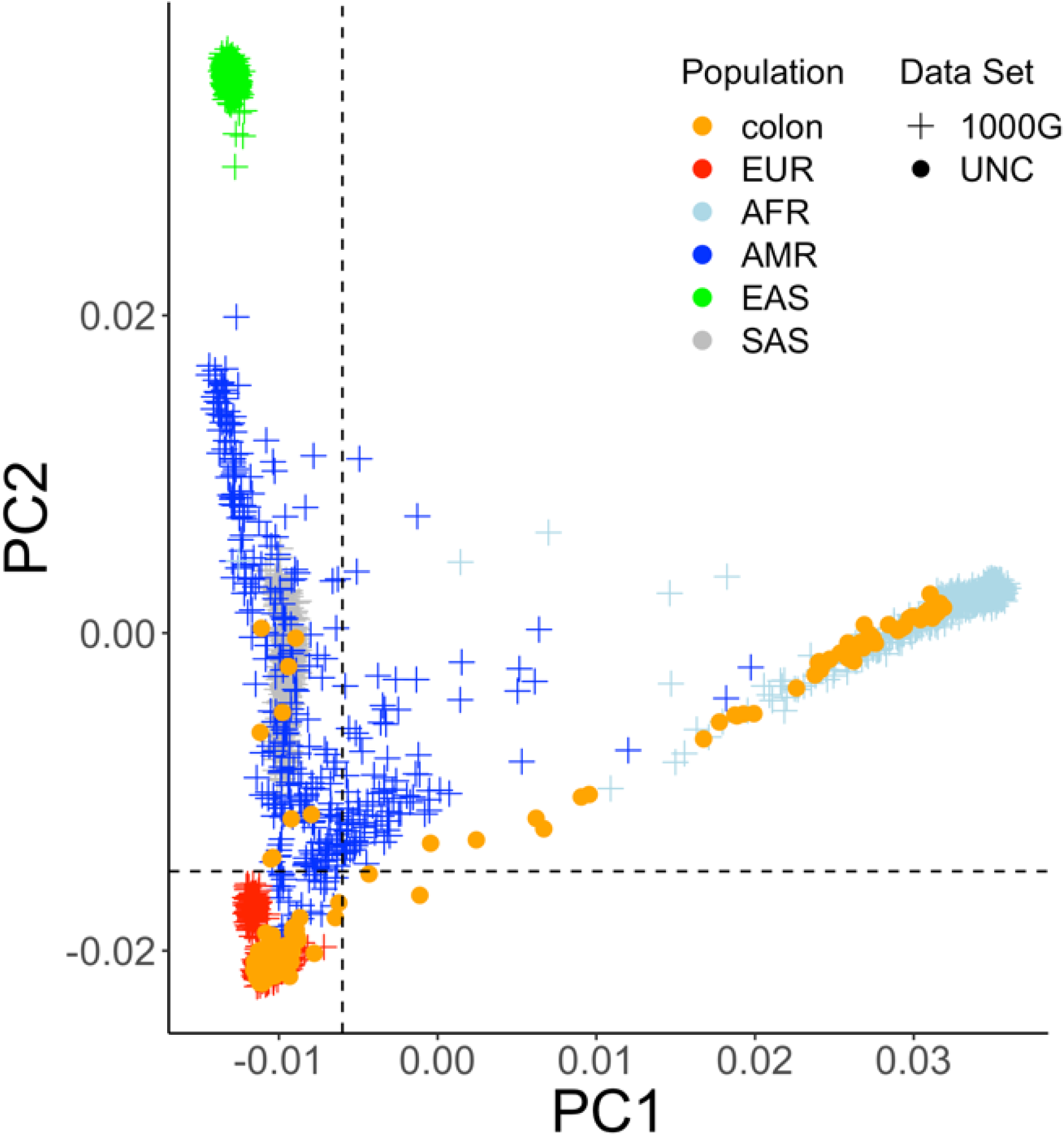
IBD patients used in this study are of primarily inferred European genetic ancestry.

**Supplementary Figure 10.**
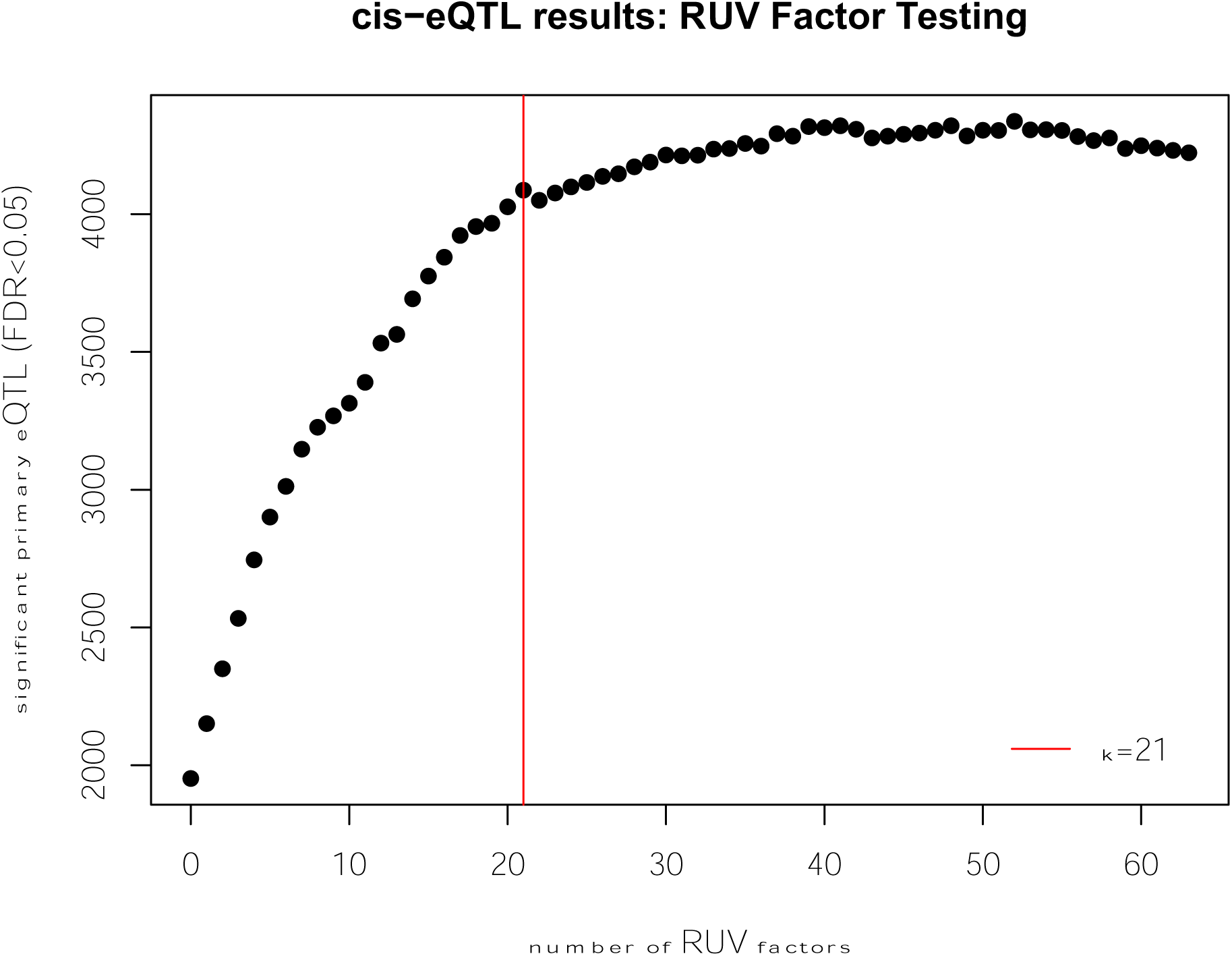
Scatter plot of the number of significant primary lead eQTL mapped using IBD tissue using increasing number of RUV factors included in eQTL model. The red vertical line indicates the point of maximum curvature detected by kneedle and the final number of RUV factors included in the eQTL model.

